# Acquisition of ampliconic sequences marks a selfish mouse *t*-haplotype

**DOI:** 10.1101/2025.01.28.635315

**Authors:** Callie M. Swanepoel, Gaojianyong Wang, Lucy Zhang, Björn Brändl, Hermann Bauer, Pavel Tsaytler, Franz-Josef Müller, Bernhard G. Herrmann, Jacob L. Mueller

## Abstract

Mendelian genetics posits equal transmission of alleles, but selfish alleles can bias the transmission of large genomic regions or entire chromosomes^1–4^. One long-standing question is how transmission bias evolves to encompass large genomic regions. *Mus musculus* (house mouse) *t*-haplotypes exhibit up to 99% transmission bias from heterozygous males^5–14^ and harbor selfish alleles^9–14^ genetically linked to large inversions spanning the proximal half of chromosome 17^15–20^. Here, by generating a high-quality, single-haplotype assembly of a *t*- haplotype, we reveal the evolution of eight large amplicons with known^11,13,21^ and candidate selfish alleles as a distinct genetic feature. Three amplicons are conserved in closely related *Mus* species, and two have known selfish alleles in the oldest inversion, implicating amplicons and an inversion drove the origins of a selfish chromosome 17 ∼3MYA. The remaining *t*- haplotype amplicons harbor gene families expressed predominantly in haploid spermatids, newly acquired retrogenes, and the most differentially expressed genes in wild-type/*t*-haplotype spermatids. Targeted deletion of a ∼1.8Mb amplicon with candidate selfish alleles on the *t*- haplotype reduces selfish transmission in heterozygous males by 3%. Notably, the evolution of selfish allele-containing amplicons and inversions on the *t*-haplotype parallels mammalian sex chromosome evolution as signatures of selfish transmission. We propose amplicon acquisition and large inversions initiate evolutionary arms races between selfish haplotypes and serve as genome-wide signatures of selfish transmission.

## Introduction

Mendelian genetics is a game of fair play, where parental alleles have an equal chance of being transmitted to offspring. Selfish alleles do not play fair; they bias their transmission to offspring at >50% frequencies. A single allele^22^ or multiple genetically linked alleles in a haplotype can cooperate to be selfish^1–4^. However, the origin and accumulation of cooperative selfish alleles into a selfish haplotype remains poorly understood.

*Mus musculus* chromosome 17 *t*-haplotypes are ideal for understanding how multiple alleles, spanning the proximal half of a chromosome, evolved to cooperate in selfish transmission^5,6,14^. Over approximately 3 million years (MY), large inversions suppressed meiotic crossing-over across a ∼40Mb region of *Mus* chromosome 17, termed *t*-complex, and gave rise to the genetically linked *t*-haplotype^15–20^. Selfish *t*-haplotype alleles cooperate in haploid spermatids of heterozygous (*wt/t^w^*^5^) male mice to bias up to ∼99% *t*-haplotype transmission to offspring^8,14,23^. The extreme transmission bias of the *t*-haplotype is countered by homozygous (*t*/*t*) mice exhibiting embryonic lethality^24^ or male sterility^25^ that prevent fixation of the *t*-haplotype in *M. musculus.* Understanding how and when the *t*-haplotype sequences evolved selfish transmission is a near century-old question hindered by the lack of a high-quality *t*-haplotype sequence assembly. Previous sequencing of *t*-haplotypes^26,27^ could not generate accurate assemblies due to the high sequence identity with the *wt* chromosome, repetitive sequences, and embryonic lethality of *t/t* mice.

To understand how the *t*-haplotype evolved selfish transmission, we generated a high-quality, single-haplotype assembly of a *t*-haplotype using homozygous (*t*/*t*) mouse embryonic stem cells. Our sequencing of the *t*-haplotype revealed three key findings. First, the *t*-haplotype acquired eight amplicons (arrays of palindromic and tandem >8kb segmental duplications with >99% identity) harboring known selfish alleles and candidate selfish alleles based on haploid spermatid expression. Second, we reconstructed the origins of the *t*-complex and how it evolved through a series of inversions and amplicon acquisition. Third, we generated the first targeted mutation on a *t*-haplotype—a ∼1.8Mb deletion of a *t*-haplotype amplicon with a candidate selfish gene family—and found reduced *t*-haplotype selfish transmission. These three findings unveil amplicons and inversions as the most prominent features of selfish haplotype evolution on *Mus* chromosome 17, strikingly similar to features of selfish alleles on mammalian sex chromosomes. Via the lens of the *t*-haplotype, we posit that amplicons and large inversions are common genomic signatures of selfish haplotypes.

## Results

### Five inversions suppress *t*-haplotype crossing over

To define the precise genomic architecture of the *t*-haplotype, we generated a haplotype-resolved assembly of the *t^w^*^5^-haplotype. *t^w5^* is ideal for delineating the evolution of the *t*- haplotype because it is a full-length haplotype (i.e., ∼ 40Mb) and exhibits all phenotypes associated with the *t*-haplotype^28–30^. To generate a *de novo*, high-quality assembly of the *t*^w5^, we used homozygous (*t^w5^/t^w5^*) genomic DNA from mouse embryonic stem cells and combined PacBio Hi-Fi, Oxford Nanopore (ONT), and BioNano long-molecule sequencing and mapping technologies. Our *t^w5^* assembly comprises 99.76Mb across three contiguous gapless scaffolds, with scaffold order and orientation validated by Hi-C contact mapping (Fig. 1, Extended Data Fig. 1A, B).

**Figure 1.**
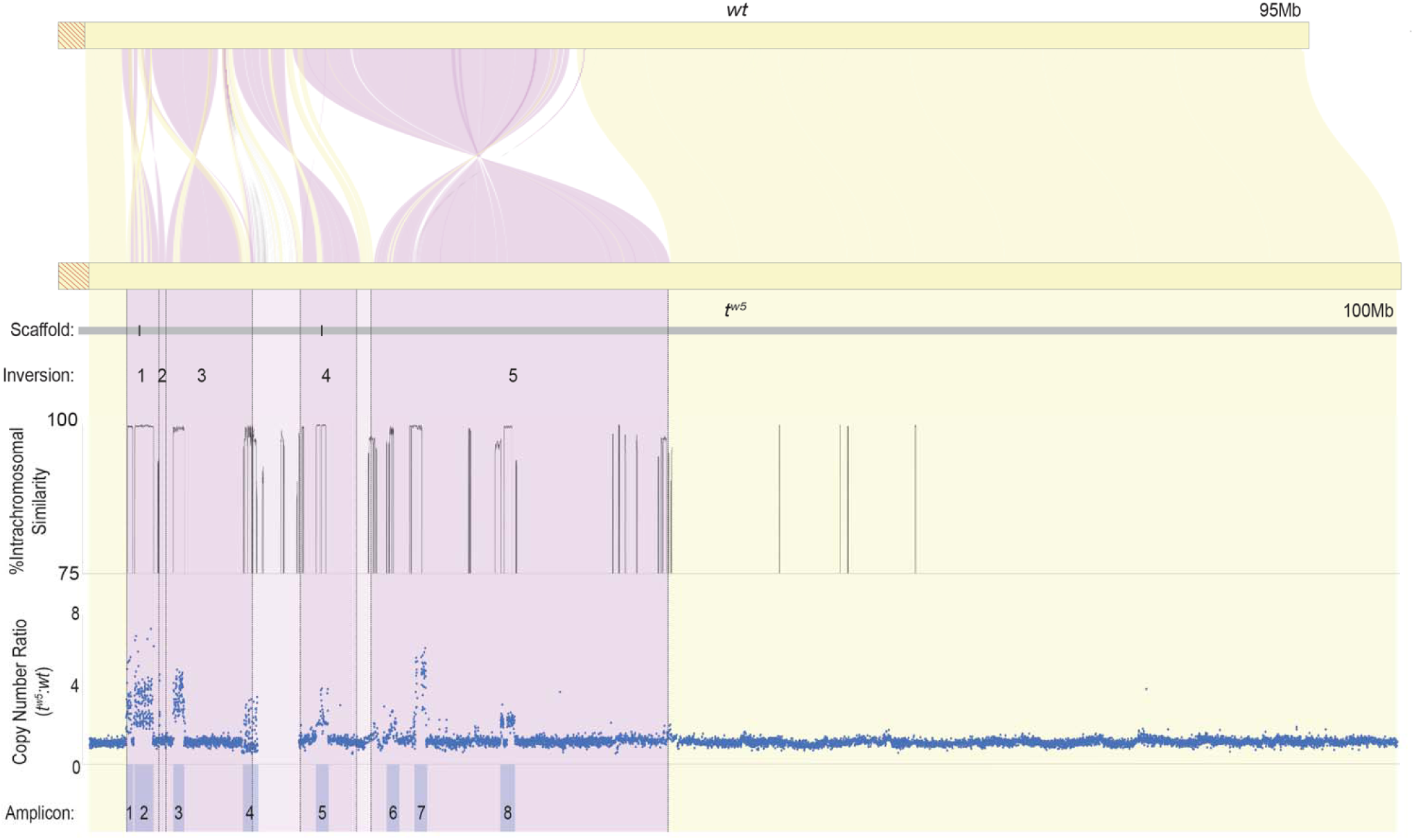
Five inversions of the mouse chromosome 17 *t^w5^* haplotype Synteny plot comparison of the mm10 reference chromosome 17 (*wt*; top) and *t^w5^* sequence assembly (bottom) with syntenic blocks in yellow and inverted regions in purple. Red diagonal lines represent the centromere. Vertical dotted lines represent inversion junctions. Three non-overlapping hybrid scaffolds (4Mb, 14Mb, and 81Mb) are represented by a grey line with black bars denoting two physical gaps between scaffolds. Five inversions are numbered between the *t^w5^* and *wt* chromosomes—a plot of intrachromosomal sequence similarity to identify the *t^w5^* ampliconic sequence. The *t^w5^* sequence was divided into 50-kb sliding windows with a 1kb step size and compared to the *t^w5^* assembly. The highest non-self-sequence similarity was plotted for each window, and all values >75% are shown. *t^w5^* copy number estimates relative to *wt* (Y-axis) are based upon read-depth analysis of whole-genome short-read sequencing data from *t^w5^*/*t^w5^*mESCs and C57BL/6J adult mice mapped to our *t^w5^* assembly (X-axis). Each blue dot represents copy number estimates in a 3kb window of unique sequence. Copy number ratios >1 represent a sequence with increased copy numbers on the *t^w5^*. We identified eight sequence blocks of increased copy number on the *t^w5^* (amplicons shaded blue; Amp1-8) relative to *wt*.

Our final *t^w5^* assembly defines the precise inversion landscape harboring *t*-haplotype selfish alleles. Previous recombination mapping of rare crossovers between the *t*-haplotype and *wt* chromosome 17 predicted four large, non-overlapping inversions within the *t*-complex^15–19^, but the precise boundaries of each inversion were not known. By comparing our *t^w5^* assembly to *M. musculus* chromosome 17 (*wt*), we confirmed four predicted inversions (1,3,4,5)^15–19^ and discovered one novel, overlapping inversion (2) (Fig. 1, Extended Data Fig. 1A, Extended Data Table 1), each inversion confirmed by PCR and Sanger sequencing of the junctions (Extended Data Fig.1C, Extended Data Table 2). We find two segments of non-inverted sequence, between inversion 3 and 4 and inversion 4 and 5, encompassing ∼2Mb of possible substrate for rare *t*-haplotype cross-over events with the *wt* allele (estimated one cross-over per 500 -1000 offspring^31^) to produce partial *t-*haplotypes^15,16,32–35^. Altogether, the *t^w5^* spans ∼41Mb of meiotic cross-over suppressed sequence, which is ∼6.8Mb (∼20%) longer than the homologous *wt* region of the C57BL/6J chromosome 17 (Extended Data Table 3).

### Eight distinct amplicons on the *t*-haplotype

Regions of the genome where meiotic crossovers are suppressed accumulate repetitive sequences and undergo single-copy sequence degeneration^36–38^. We asked whether these repetitive sequence types account for the ∼6.8Mb difference in *t^w5^* total sequence relative to *wt*. We find the *t^w5^* has ∼11.7Mb more repetitive sequence as compared to *wt*, including 6.5Mb of amplicons across eight regions (Amp1-8) with >99% intrachromosomal sequence identity (Fig. 1, Extended Data Fig. 2A and B, Extended Data Fig. 3, Extended Data Table 4), 2.8Mb of satellite sequence (Extended Data Fig. 2C and Extended Data Table 3), and 2.1Mb of transposable elements (Extended Data Table 3). Since the *t^w5^* has 11.7Mb more repetitive sequence than *wt,* this suggests non-repetitive sequences degenerated on the *t^w5^* to explain the overall ∼6.8Mb difference in *t^w5^*total sequence length. Indeed, the *t^w5^* has 3.8Mb of single-copy sequence with low alignment coverage relative to *wt* (Extended Data Fig. 1D and Extended Data Table 3), indicating loss or highly diverged sequence. Thus, the *t^w5^* sequence appears to have accumulated repetitive sequences and degenerated single-copy sequence, consistent with known patterns of chromosomal regions with suppressed meiotic crossing over^37,38^.

Since amplicons are the most distinct sequence on the *t*-haplotype relative to *wt*, we further characterized their structure and copy number. To quantify *t^w5^*sequence amplification, we determined each amplicon’s relative copy number and structural arrangement on the *t^w5^* versus *wt* (Fig. 1 and Extended Data Fig. 2B). We found all eight *t*-haplotype amplicons have higher copy number relative to *wt*, with three regions amplified on both the *t^w5^* and *wt,* and five regions uniquely amplified on *t^w5^* (Fig. 1, Fig. 2 and Extended Data Fig. 3). The three shared amplicons between *t^w5^* and *wt* suggest a common ancestor. In contrast, the *t*-haplotype-specific amplicons could have been gained on the *t*-haplotype or lost from the *wt* chromosome.

**Figure 2.**
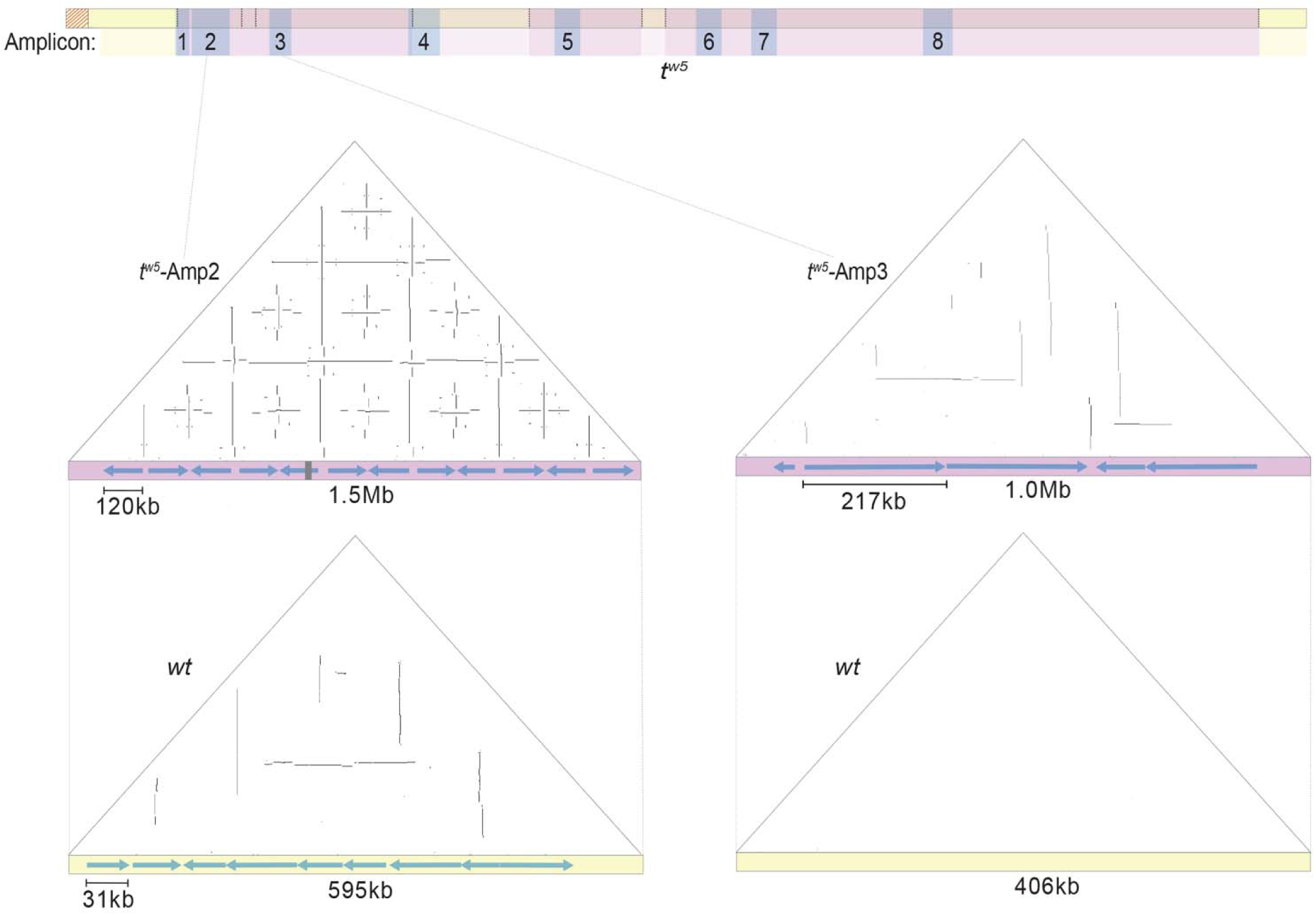
*t^w5^* sequence is amplified relative to *wt* A schematic of the *t^w5^* with amplicons shaded blue and dotted lines representing inversion junctions. Below are self-symmetry dot-plots highlighting palindromic (vertical lines) and tandem (horizontal lines) segmental duplications within *t^w5^*-Amp2 (top left) and the orthologous *wt* region (bottom left) that is amplified to a lower copy number. Each dot represents a 100bp window of 100% nucleotide identity. The X-axis is the sequence, 5’ to 3’, and blue arrows represent the positions and orientations of each >8kb, >99% identical amplicon. The gray bar denotes a physical gap in the assembly. Dot-plots on the right compare *t^w5^*-Amp3 (top), uniquely amplified on the *t^w5^*, to the orthologous *wt* region (bottom).

### Conserved amplicons in the oldest inversion point to the origin of the *t*-haplotype

To reconstruct the evolutionary history of *t*-haplotype amplicons, we asked whether amplicons evolved on chromosome 17 in closely related *Mus* species. We estimated chromosome 17 copy number in *M. caroli*, *M. spretus,* and *M. spicilegus* (Fig. 3A and Extended Data Fig. 4A). We found five *t*-haplotype amplicons are acquired recently only on the *t*- haplotype, as they are not amplified in any of the four *Mus* species examined. Relative to *M. caroli*, we find the three amplicons shared between *M. musculus t*-haplotype and *wt* are conserved in *M. spretus* and *M. spicilegus* (Fig. 3A and B and Extended Data Fig. 4A and C).

**Figure 3.**
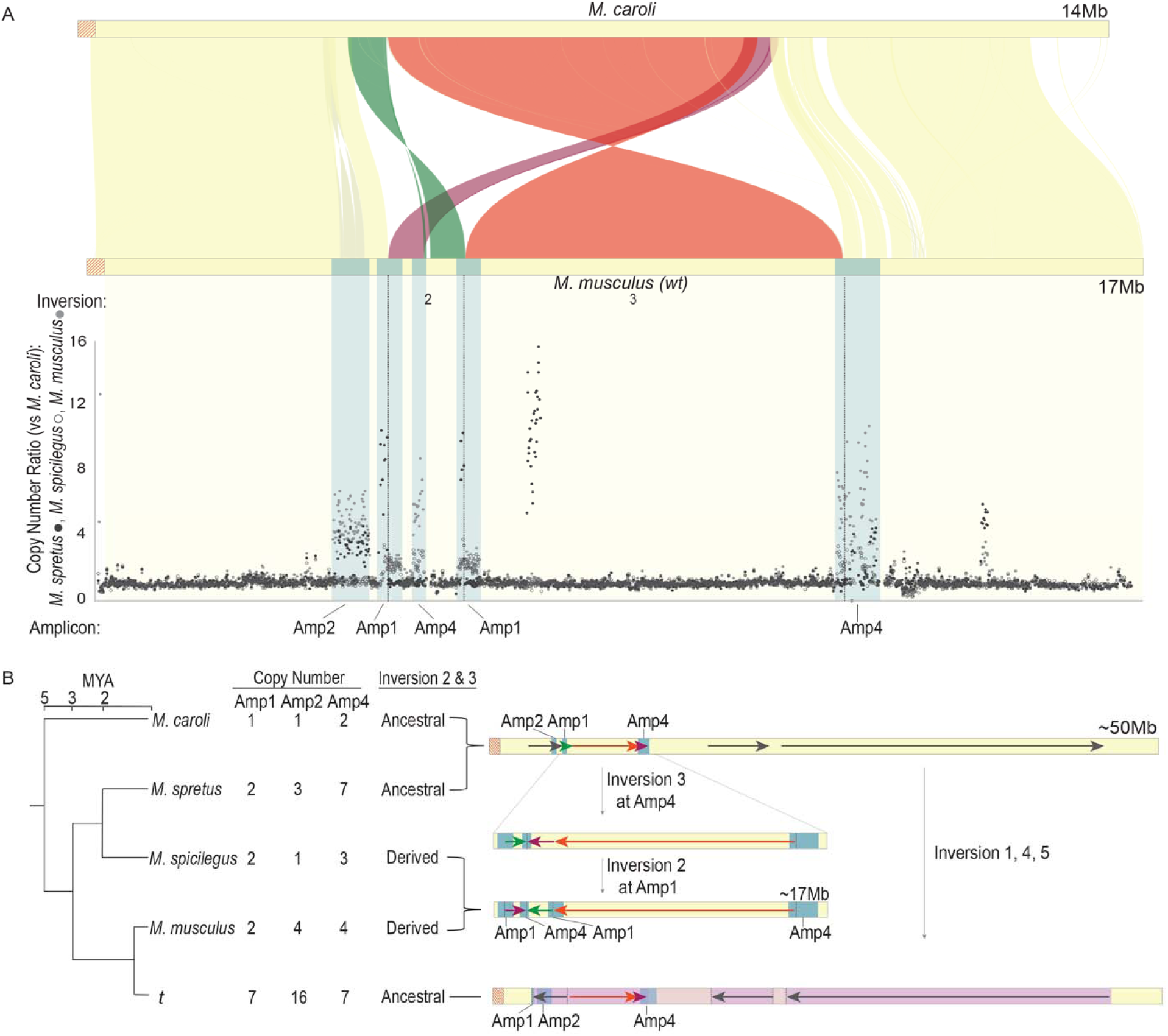
Conserved amplification genetically linked to the *t*-complex oldest inversion **A**. (Top) Synteny plot comparison of the first 14Mb of *M. caroli* chr 17 and first ∼17Mb of *M. musculus* chr17. Syntenic blocks are shaded in yellow, and inverted regions are shaded in green, orange, or maroon. Grey shading indicates amplified sequence (Amp2) on M*. musculus* chr17. Blue shading along the *M. musculus* chromosome represents regions of conserved amplification. Dotted lines represent inversion junctions. (Bottom) Copy number estimates across the first ∼17Mb of *M. spretus*, *M. musculus,* and *M. spicilegus* chr17 relative to *M. caroli* (Y-axis). Each dot represents a copy number estimate across a 3kb window of a unique sequence. Conserved amplified regions are labeled below the X-axis. **B.** (Left) A schematic phylogenetic tree of *Mus* chr 17 with the inversion 2 and 3 orientation (ancestral or derived) and copy number estimates for conserved amplicons (Amp1, 2, and 4) next to each species. (Right) A model of inversions 3 and 2 from the ancestral state to derived (*M. musculus wt* and *M. spicilegus*) and inversions 1, 4, and 5 from the ancestral state to the *t* (bottom). Black dotted arrows denote inversion junctions, and blue shading indicates amplified regions. Green, maroon and orange horizontal arrows correspond to inverted regions from panel A, and grey arrows correspond to inverted regions on the *t*. Along the *t* chromosome, dark purple shading represents inverted regions, and light purple shading represents non-inverted regions between *t* and *wt*. Red diagonal lines represent the centromere.

Notably, two of the three conserved amplicons (Amp1 and Amp4) harbor *Tagap* and *Smok2* gene families, which have been previously implicated to have selfish *t*-haplotype alleles^11,13^ (Fig. 4A, Extended Data Fig. 4A and C and Extended Data Table 4). The third conserved amplicon (Amp2) harbors *Dynlt1* and *Tmem181a* gene families that are unexplored for their contribution to selfish transmission (Fig. 4A, Extended Data Fig. 4A and C and Extended Data Table 4).

**Figure 4.**
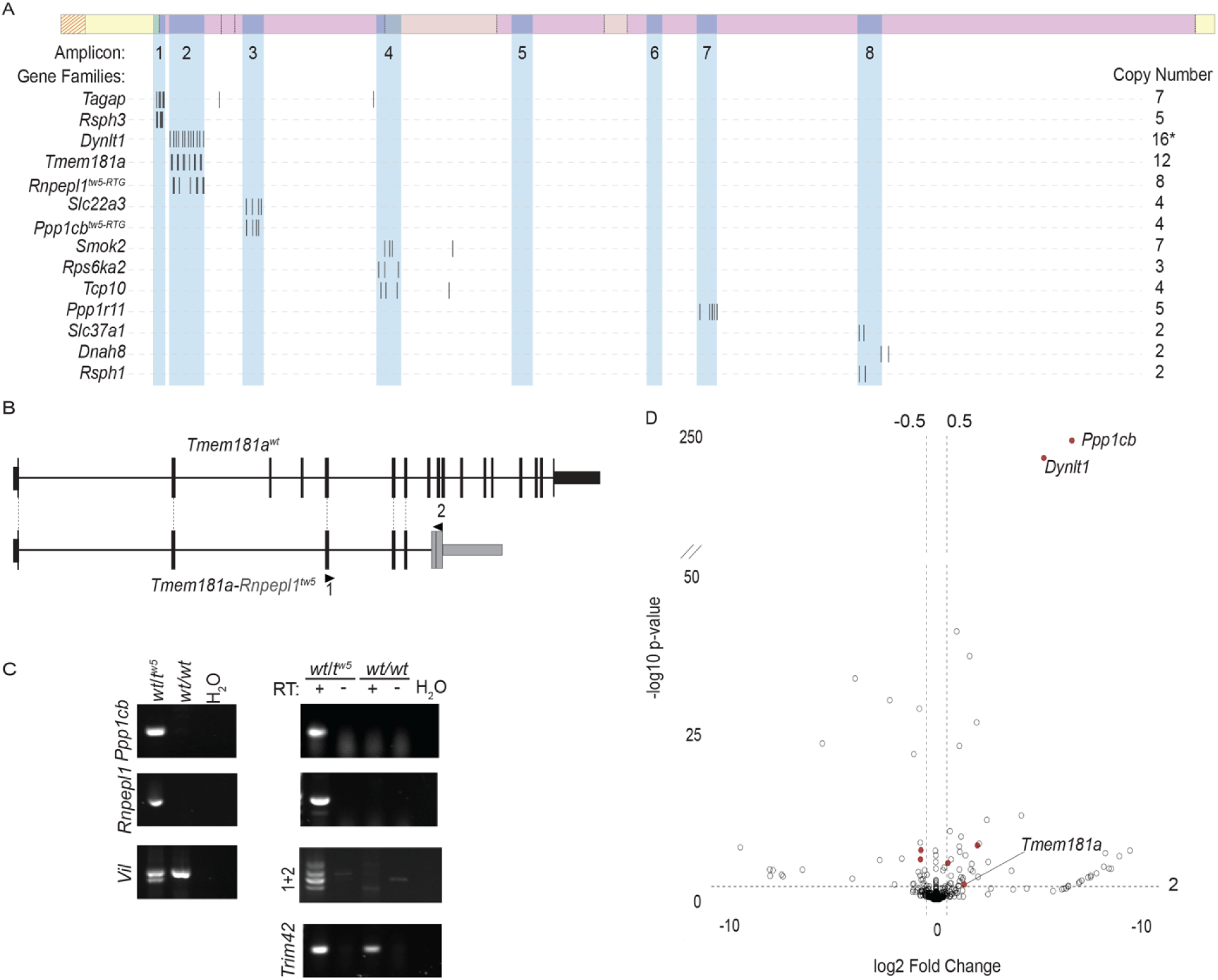
*t^w5^* amplicons with known and candidate gene families involved in selfish transmission **A.** The *t^w5^* with ampliconic regions shaded in blue and numbered below. Below are the positions of ampliconic gene families with haploid spermatid expression (black ticks) and their copy numbers. **B.** Our gene annotation schematic representation of *Tmem181a-Rnpepl1^tw5^* fusion gene structure as compared to *Tmem181a^wt^*. Black vertical bars represent exons, and horizontal lines represent introns. Dotted lines connect orthologous exons between genes. The regions of *Rnpepl1* that retrotransposed to the *t^w5^* include part of exon 10, exon 11, and the 3’ UTR and are shaded in gray. Arrows with numbers denote positions of RT-PCR primers used in panelC **C.** PCR (left) validation of *Ppp1cb* and *Rnpepl1* retrotransposition from *t^w5^*but not *wt* genomic DNA. *Vil* is an internal positive control to genotype *t^w5^* (two-bands) vs *wt* (single-band). RT-PCR (right) validation of retrotransposed *Ppp1cb*, *Rnpepl1,* and *Tmem181a-Rnpepl1* testis expression. Four different bands amplified across the *Tmem181a-Rnpepl1* fusion junction were cloned and sequenced for validation. *Trim42* is a positive control for spermatid-expressed genes. + and - denote RT positive and no RT controls. **D.** Volcano plot of differentially expressed genes (-log10 p-value > 0.01; log2 fold change > 0.5) in *t^w5^/wt* versus *wt/wt* haploid spermatids. Filled red dots represent *t^w5^* ampliconic gene families with differential expression. Diagonal lines represent a break in the y-axis.

While the selfish *t*-haplotype is only observed in *M. musculus*, the conserved amplification of known gene families with selfish alleles suggests selfish transmission initiated in a common ancestor of *M. spretus* and *M. musculus*.

If conserved amplicons initiated selfish transmission, we predict they would be associated with the most ancestral *t*-complex inversions. To identify the oldest inversion, we compared the order and orientation of chromosome 17 assemblies for *M. spretus*, *M. spicilegus*, *M. musculus,* and the *t^w5^* relative to *M. caroli* (Fig. 3 and Extended Data Fig. 4B). We find inversions 1, 4, and 5 are unique to *t^w5^*, supporting their recent origin on the *t*-haplotype (Fig. 3B and Extended Data Fig. 4B and E). Inversions 2 and 3 are derived (inverted) in *M. spicilegus* and *M. musculus* and ancestral (non-inverted) on the *t*^w5^ relative to *M. caroli* (Fig. 3 and Extended Data Fig. 4B-E). Thus, inversions 2 and 3 evolved on an independent haplotype from inversions 1, 4, and 5 (Fig. 3B and Extended Data Fig. 4E). This finding is consistent with previously observed crossing over in the region spanning inversions 2 and 3 between a *t*- haplotype and *M. spretus* chromosome 17, but not between *M. musculus* (*wt*) and *M. spretus* chromosome 17^17^. To achieve the order and orientation of inversions 2 and 3, inversion 3 must precede inversion 2, which we confirmed via PCR of the derived (*M. spicilegus* and *M. musculus*) and ancestral (*M. spretus* and *t^w5^*) inversion junctions (Extended Data Fig. 4D). Based on these findings, we conclude that inversion 3 is the most ancestral inversion, followed by inversion 2. Notably, *Smok2* and *Tagap*, gene families with known selfish *t*-haplotype alleles, map to the breakpoints of inversion 3 and inversion 2. Our finding of conserved amplification of *Smok2* and *Tagap* gene families being linked to the oldest inversions (2 and 3) suggests they initiated the evolution of selfish transmission.

### *t-*haplotype amplicons harbor known and new candidate selfish alleles

*Smok2* and *Tagap* are among five genes previously implicated to have selfish *t*- haplotype alleles (*Fgd2*, *Nme3*, *Tiam2, Tagap1*, and *Smok^Tcr^)*^9–11,13^ based on sufficiency studies with partial *t*-haplotypes. Of these five genes, *Smok2* and *Tagap* are the only gene families within *t^w5^* amplicons (Fig. 4A and Extended Data Table 4). Nonetheless, the discovery of known selfish alleles within *t^w5^* amplicons suggests other amplicon gene families have alleles contributing to *t*-haplotype selfish transmission.

We asked whether the remaining *t*^w5^ amplicons are potential regions for selfish allele evolution based on the presence of genes with haploid spermatid expression, the cell type where known selfish alleles act molecularly^39^. To identify new candidate *t*-haplotype amplicon gene families contributing to selfish transmission, we gene annotated each amplicon and examined haploid spermatid gene expression. We annotated the amplicons for gene orthologs to *wt* chromosome 17 and newly acquired genes within *t^w5^*amplicons. We identified 14 *t^w5^*amplicon gene families with *wt* orthologs and *wt/t^w5^* haploid spermatid expression (Fig. 4A and D and Extended Data Table 4). To identify newly acquired *t^w5^*genes, we generated a *de novo t^w5^* gene annotation and revealed four *t^w5^* specific retrotransposed genes (*t^w5^*-RTG) (Extended Data Fig. 5), two of which were previously proposed *t*-haplotype retrogenes, but their copy number and gene structure were not characterized^40^. Two *t^w5^* retrogenes, *Rnpepl1^tw5-RTG^*and *Ppp1cb^tw5-RTG^*, are within amplicons and have *wt/t^w5^* haploid spermatid expression (Fig. 4A and C and Extended Data Fig. 5). *Rnpepl1^tw5-RTG^*copies include only the last two exons and 3’UTR of the *Rnpepl1* progenitor gene and some copies form multiple fusion transcripts with *Tmem181a* (Fig. 5B and C). PCR cloning and Sanger sequencing of *Tmem181a-Rnpepl*1^t*w5-RTG*^ fusion transcripts reveal alternate splicing within *Rnpepl*1*^tw5-RTG^* and no apparent open reading frame, suggesting *Rnpepl*1*^tw5-RTG^* leads to loss of some *Tmem181a* functional copies. In contrast to the gene fusion associated with *Smok2*^13^, the *Tmem181a-Rnpepl*1*^tw5-RTG^* gene fusion appears to be disruptive. Since haploid spermatids are the cell type where selfish *t*-haplotype genes act, the spermatid expression of amplicon gene families and associated loss or gain of function mutations on the *t*-haplotype make them exciting candidate selfish alleles.

**Figure 5.**
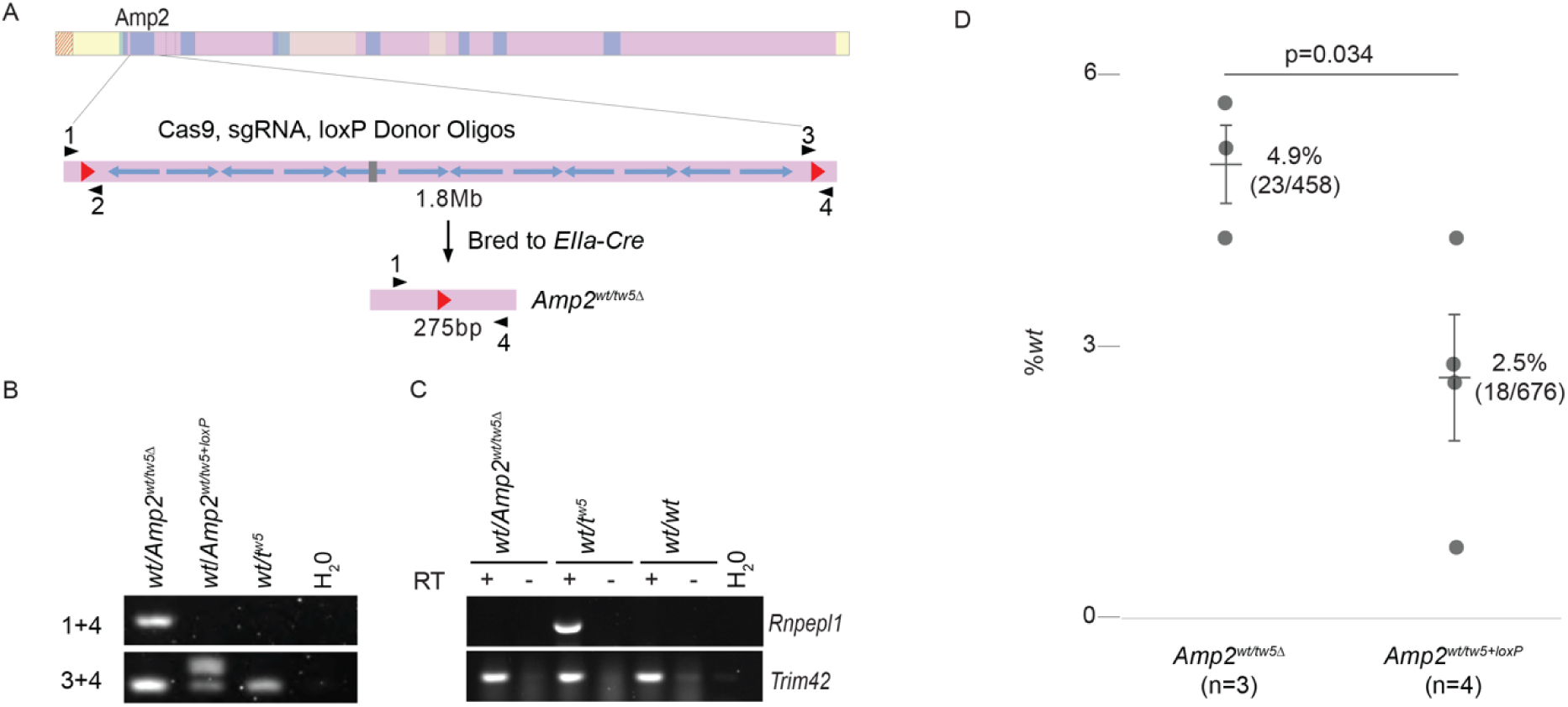
Deletion of *t^w5^*-Amp2 reduces selfish transmission **A.** A schematic representation of the *t^w5^*with ampliconic regions shaded in blue. Below is a schematic of the *t^w5^*-Amp2 region targeted for deletion via sequential integration of loxP sites (red arrows) using sgRNA-guided Cas9 and germline deleted by crossing to mice carrying an *Ella*-Cre transgene. Black arrows with numbers denote primer positions for genotyping. **B.** PCR validation of the *Amp2 ^tw5Δ^* from genomic DNA. **C.** RT-PCR validation of the *Amp2 ^tw5Δ^* using primers specific to the *Rnpepl1^tw5^*retrogene within *t^w5^*-Amp2. **D.** *Amp2^+/tw5Δ^* (n=3), *Amp2^+/tw5-loxP^* (n=4) males were mated against WT females, and progeny were genotyped for *Amp2 ^tw5Δ^* and *t^w5^* vs *wt* alleles. The average percent *wt* transmission (horizontal lines) and total number of *wt* alleles/total progeny genotyped in parentheses are shown. Vertical lines represent the standard error for each group. The p-values were calculated using a generalized linear mixed model. Each dot represents the percent *wt* alleles transmitted from a single stud male.

### Differential expression of *t*-haplotype amplicon gene families in haploid spermatids

Previously implicated *t*-haplotype selfish alleles are differentially expressed in *wt*/*t^w5^* versus *wt/wt* testes^9–12^. Thus, if *t^w5^* amplicon gene families have a role in selfish transmission, they may be differentially expressed in *wt*/*t^w5^*versus *wt/wt* haploid spermatids. We find *Ppp1cb^tw5-RTG^* and *Dynlt1*, both within *t^w5^* amplicons, are the top two most over-expressed genes relative to *wt* (Fig. 4D and Extended Data Fig. 6). *Ppp1cb^tw5-RTG^* arose from a chromosome 5 progenitor copy of *Ppp1cb*, and acquired spermatid expression since *Ppp1cb* is only expressed in spermatogonia and spermatocytes^41^ (Fig. 4D and Extended Data Fig. 6). Whether the *Ppp1cb^tw5-RTG^*acquired novel spermatid function remains to be tested. *Dynlt1* is the most amplified gene family on *t^w5^* and is within conserved Amp2 (Fig. 2, Fig. 3, Fig. 4A and Extended Data Table 4). Because there is an assembly gap in the *t^w5^*-Amp2 region where the *Dynlt1* gene family resides, we used digital droplet PCR (ddPCR) to determine that *Dynlt1^tw5^* has 16 copies and *Dynlt1^wt^* has four copies (Extended Data Fig. 7). The ∼5-fold increase in *Dynlt1* RNA expression in *wt*/*t^w5^* spermatids is concordant with a 4X increase in *Dynlt1^tw5^*DNA copy number. Similar to previously identified *t*-haplotype selfish alleles^9,11^, differential expression of *Ppp1cb* and *Dynlt1 t^w5^* amplicon gene families indicates their potential contribution to selfish transmission.

### *A t*-haplotype amplicon encoding differentially expressed genes contributes to selfish transmission

To address the functional contribution of *t*-haplotype amplicons to selfish transmission, we tested the necessity of *t^w5^*-Amp2, harboring candidate selfish gene families: *Dynlt1*, *Rnpepl1^tw5-RTG^*, *Tmem181a,* and *Tmem181a- Rnpepl1^tw5-RTG^* (Fig. 5). We selected *t^w5^*-Amp2 because it is one of the three conserved amplicons in *M. spretus* (Fig. 3 and Extended Data Fig. 4), is the most amplified region on the *t^w5^* (∼16 copies) (Fig. 2, Fig. 4A and Extended Data Fig. 7), harbors the second most overexpressed gene family (*Dynlt1*) in haploid spermatids from *wt/t^w5^* versus *wt/wt* males (Fig. 4D and Extended Data Fig. 6), and *Dynlt1* encodes dynein protein, a protein family known to be involved in sperm motility and sperm axoneme formation^2,42,43^. We generated a targeted ∼1.8Mb deletion of *t^w5^*-Amp2 and assessed if selfish transmission of *t^w5^* from *Amp2^wt/tw5Δ^* males is reduced. We observe decreased *t^w5^* transmission from ∼98% to ∼95% in *Amp2^wt/tw5Δ^* versus *Amp2^wt/tw5-loxP^* (p=0.034) males. By comparison, the deletion of the orthologous *wt*-Amp2 region (*Amp2^wtΔ^*) does not impact chromosome 17 transmission (Extended Data Fig. 8, p= 0.664), suggesting a *t*-haplotype specific allele within *t^w5^*-Amp2 contributes to selfish transmission. We consider *Dynlt1* the most likely gene family contributing to *t^w5^*-Amp2-mediated selfish transmission based on its evolution, expression, and predicted protein function. We cannot exclude the possibility of *Tmem181a* contributing to selfish transmission since it is upregulated in haploid spermatids from *wt/t^w5^* versus *wt/wt* (Figure 4D). Since *Rnpepl1^tw5-RTG^* and the *Tmem181a-Rnpepl1^tw5-RTG^* fusion transcripts lack open reading frames, they are less likely contributors to selfish transmission. Regardless of which gene family contributes to selfish transmission, deletion of *t^w5^*-Amp2 supports the contribution of amplicons to *t* selfish transmission.

## Discussion

Our study of a *t*-haplotype revealed amplicons with known and candidate selfish *t*- haplotype alleles, suggesting amplicon acquisition is a key genomic signature of selfish haplotype evolution. Most *t*-haplotype amplicons harbor spermatid-expressed genes, thus potentially being selfish. Indeed, spermatid-expressed ampliconic genes arose with the initiation and evolution of the *t*-complex, and deletion of *t^w5^*-Amp2 reduces selfish transmission. The 3% contribution of *t^w5^*-Amp2 can significantly impact *t*-haplotype transmission over evolutionary time. The *t^w5^*-Amp2 region likely cooperates with other selfish *t*-haplotype alleles, culminating in extreme selfish *t*-haplotype transmission. This cooperation is supported by *t*- haplotype alleles in inversion 1 (where *t^w5^*-Amp2 is located) and inversion 5 additively influencing selfish transmission^44^, and partial *t*-haplotypes without inversion 1 reducing selfish transmission^8,14^. Our studies lay the foundation for future functional studies to address the cooperation of known and candidate selfish *t*-haplotype alleles.

Based on the evolutionary history of the *t*-complex, we propose inversions, selfish alleles, and amplification initially arose on the ancestral *wt* haplotype and were countered by subsequent inversions and amplicon acquisition on the *t*-haplotype (Fig. 3 and Extended Data Fig. 4). We speculate that an evolutionary arms race for selfish transmission driven by inversions and amplicon acquisition continually shapes *Mus* chromosome 17 evolution, leaving the intriguing question of whether independent inversions or amplicons evolved in other *M. musculus t*-haplotypes or *Mus* species. Our high-quality *t*-haplotype sequence and cross-species comparisons provide initial insight into this question (*e.g.*, Extended Data Fig. 4 showing an independently acquired amplicon in *M. spretus* and additional inversion in *M. spicilegus*) and are a necessary starting point for future comprehensive sequence comparisons.

Evolution of the *t-*haplotype shares striking similarities with mammalian sex chromosome evolution. The mammalian X and Y chromosomes diverge by suppression of crossing over (likely via inversions)^37^, independently acquire multiple amplicons^45–48^, and exhibit selfish transmission due to an evolutionary arms race between spermatid-expressed ampliconic gene families^49,50^. These combined features are considered unique to mammalian sex chromosomes; however, our study finds the *t*-haplotype shares these genomic features despite >150 million years difference in age. Unlike the mammalian X and Y chromosomes, recent *t*-haplotype evolution and sequence similarity to *wt* allow a more precise reconstruction of its origin.

Evolutionary parallels between mammalian sex chromosomes and the *t*-haplotype suggest large inversions and amplicon acquisition may be genome-wide features of past or ongoing evolutionary arms races for selfish transmission.

Beyond the *t*-complex and mammalian sex chromosomes, do other large inversions harbor selfish amplicons? Selfish amplicons within large inversions are historically difficult to detect due to challenges in (**1**) resolving large inversions^51^, (**2**) identifying amplicons^52–54^, and (**3**) tracking transmission biases not associated with overt phenotypes (*e.g.*, sex for X and Y chromosomes, and short tails for the *t*-haplotype). High-quality human and mouse genome assemblies reveal 49 and 16 amplicons with spermatid-expressed gene families, respectively, representing potential regions of ongoing or past selfish transmission (Extended Data Fig. 9). At least one human autosomal amplicon with spermatid-expressed genes is within a known polymorphic inversion under selection, like the *t*-haplotype^55^ (Extended Data Fig. 9). Amplicon-containing inverted haplotypes with selfish transmission may thus be more common than previously thought. Amplicons can also harbor non-germline expressed genes (*e.g.*, MHC locus^56^ and olfactory receptors on the *t*-haplotype) (Extended Data Fig. 9), indicating amplicons may evolve in response to degeneration of an inverted chromosomal region, such as to facilitate recombination to repair deleterious alleles or increase gene expression by having multiple copies. Our high-quality assembly and analyses of a mouse *t*-haplotype are a road map for studying the roles of amplicons in additional haplotypes with selfish transmission across species.

We propose a model of how selfish amplicons and large inversions shape chromosome evolution (Fig. 6). Selfish amplicons within an inverted haplotype (H1) can become fixed in the population or engage in an evolutionary arms race for selfish transmission with the ancestral haplotype (A) or new haplotypes (H2), as with the *t*-haplotype or sex chromosomes. Whether the outcome is fixation or an ongoing arms race, amplicons and large inversions reshape chromosome evolution. Over evolutionary time, the fair play of many Mendelian genes may be impacted by episodes of selfish haplotype foul play.

**Figure 6.**
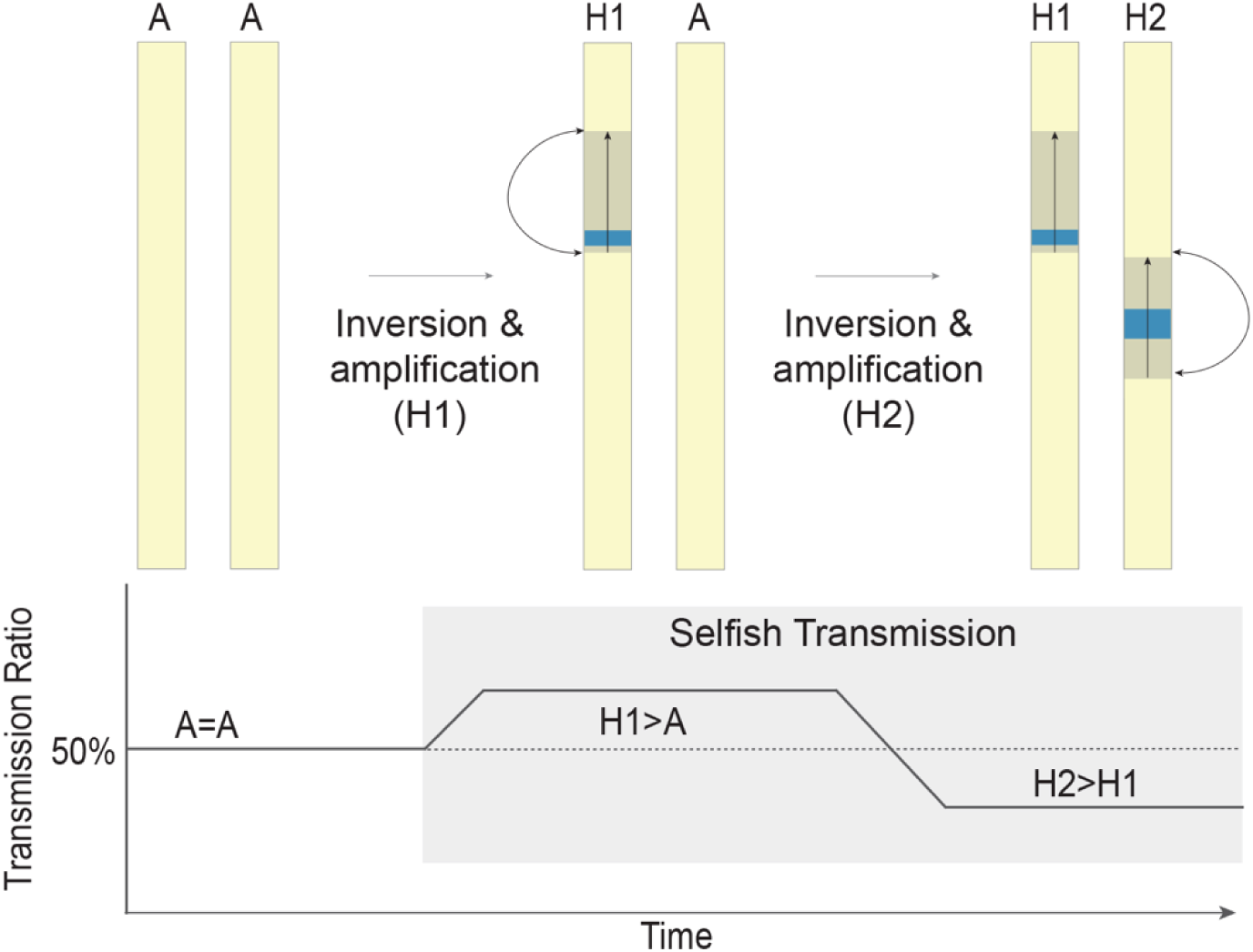
A model of a selfish transmission arms race A model of homologous chromosomes that undergo inversions (shaded gray with black arrows) and amplicon acquisition (blue) to gain selfish transmission. “A” represents the ancestral chromosome, and “H1” and “H2” represent independent haplotypes competing for selfish transmission. The graph below models the change in transmission ratio with the acquisition of inversions and amplicons on each haplotype over evolutionary time.

## Methods

### t^w5^ sequencing

The mouse *t^w5^* haplotype was sequenced using a combination of PacBio HiFi, Oxford Nanopore, and BioNano optical mapping. All sequencing data was generated using high molecular weight (HMW) DNA isolated from *t^w5^/t^w5^* mouse embryonic stem cells (mESCs) on a C57BL/6 genetic background.

<u>PacBio HiFi:</u> PacBio HiFi sequencing was conducted with HMW DNA isolated from 2- 4x10^6^ *t^w5^*/*t^w5^* mouse embryonic stem cells (mESCs) per replicate using Analytik Jena’s smart-m-Prep-kit. 2-5µg of HMW DNA was sheared to 20-30kb fragments using the MegaRuptor (Speed 31/32, T=120min) and subjected to 1x AmpurePB cleanup. PacBio circular consensus/HiFi-libraries were prepared from 4µg of fragmented DNA following PacBio’s SMRTbell Express Template Preparation protocol. Size selection was performed using a BluePippin with an HSLF chip, and DNA between 10-50kb was eluted and isolated using AmpurePB beads. The resulting libraries range between 20 and 28kb fragments (FragmentAnalyzer) and were loaded on four PacBio SequelII SMRTcells using adaptive loading. After 30h of data acquisition between 39 and 142Gb (38.89, 141,72, 104.41 and 133.3Gb) of unique Molecular Data was yielded per sample (260.91, 541.38, 539.61, 581.01Gb Total Bases).

<u>Oxford Nanopore</u>: Nanopore ultra-long DNA sequencing was performed using UHMW DNA isolated from the Nanobind CBB Big DNA kit (Circulomics, cat. # NB-900-001-01). Briefly, three aliquots of 6x10^6^ *t^w5^*/*t^w5^*mouse embryonic stem cells (mESCs) were subjected to DNA extraction following the manufacturer’s protocol. The eluates were then quantified using the Qubit dsDNA BR kit, and ∼40µg of UHMW DNA was subjected to the Nanobind UL Library Prep kit (Circulomics, cat. # NB-900-601-01) in conjunction with ONT’s Ultra-Long DNA Sequencing kit (Oxford Nanopore, cat. # SQK-ULK001).

Tagmentation was performed using 6 µl of FRA enzyme and subsequent ligation of the RAP-F sequencing adapter for 1h at room temperature. The final sequencing libraries were loaded onto R9.4.1 MinION flow cells (Oxford Nanopore, FLO-MIN106) for N50 fragment length estimation and then processed on two R9.4.1 PromethION flow cells (Oxford Nanopore, FLO-PRO002) for 72h. A nuclease flush (Oxford Nanopore, EXP- WSH003) was performed every 16h according to the manufacturer’s protocol, and a fresh sequencing library was loaded. A total of ∼121Gb (∼48x coverage) basecalled bases with N50 read length of ∼90kb were obtained and further subjected to genome assembly.

<u>BioNano Optical Mapping:</u> Ultra-High Molecular Weight DNA was extracted from frozen cells using the “Bionano Prep SP Frozen Cell Pellet DNA Isolation Protocol v2”. The long DNA molecules were labeled with the DLE-1 enzyme according to the “Bionano Prep Direct Label and Stain (DLS) Protocol.” The labeled molecules were loaded into a single flowcell of a Saphyr Chip G1.2. The chip was run on the Bionano Saphyr Optical Mapping Instrument twice, collecting the images of 955.75 Gbp molecules greater than 150kb with a minimum of 9 labels. These data were assembled using Bionano Solve v.3.6.1 software and optArguments_nonhaplotype_DLE1_saphyr.xml parameters into 739 genome maps with an N50 of 99.13Mb and a total length of 2725.521Mb.

### t^w5^ chromosome-level assembly

A chromosome-level assembly was generated from PacBio HiFi, ONT, and Bionano long molecule technologies. Briefly, Guppy v6.3.7 (Oxford Nanopore Technologies, UK) was used for basecalling ONT raw reads, utilizing the configuration file dna_r9.4.1_450bps_hac.cfg. The generated sequences are named ONT data. To obtain PacBio HiFi consensus reads, we followed the recommended workflow of DeepConsensus^57^, where the subreads of each run were partitioned into 500bp segments. We obtained the draft consensus sequences using ccs v6.3.0 (PacificBiosciences, USA) and aligned the subreads to the draft consensus sequences using actc v0.3.1 (PacificBiosciences, USA) for each segment of partitioned data. We then applied DeepConsensus^57^ v1.0.0 to the draft consensus sequences and aligned the subreads of each 500bp segment to generate more accurate consensus sequences. The generated accurate consensus sequences from DeepConsensus^57^ are named HiFi data. We used Verkko^58^ v1.3, using both HiFi and ONT data with default parameters to generate a de novo assembly.

Assembled sequence contigs were placed onto BioNano optical map scaffolds using Bionano Solve v.3.6.1 software and default hybridScaffold_DLE1_config parameters, resolving conflicts in both the optical map and sequence assemblies to generate large contigs. Three non-overlapping contigs comprising the *t^w5^* chromosome were identified using BLAST^59^ to compare all contigs to the proximal 40Mb of mm10 chr17 that was repeat masked using default parameters of RepeatMasker^60^. The order and orientation of each contig was determined based upon shared repeat content at the 3’ and 5’ ends of each scaffold and independently validated using Hi-C scaffolding (Extended Data Fig. 1A). While there are no sequence gaps within contigs, there are two remaining physical gaps between contigs represented by 25kb of N’s.

### Validation of t^w5^ assembly

To independently validate the *t^w5^* assembly, we use PacBio CLR to generate a de novo assembly and Hi-C to confirm the order and orientation of the contigs (Extended Data Fig. 1A).

<u>PacBio CLR:</u> PacBio CLR sequencing was performed with 15µg of HMW DNA from *t^w5^*/*t^w5^* mESCs. For library preparation, HMW DNA was sheared to a target size of >30kb using the MegaRuptor (Speed 3, t=159m49s) and cleaned up using AmpurePB beads.

PacBio CLR libraries were prepared from 4µg of fragmented DNA following PacBio’s SMRTbell Express Template Preparation protocol. Size selection was performed using a BluePippin with a HSLF chip, and 30kb DNA fragments were eluted and cleaned up using AmpurePB beads. The final library fragment size was 43kb, according to Fragment Analyzer. Two flowcells were run with 15h acquisition time on the PacBio SequelII, yielding 78 and 142Gb unique data (98 and 165Gb total data).

<u>Hi-C:</u> Hi-C was performed using 2x10^6^ *t^w5^/t^w5^* mESCs. The cells were collected by trypsinization, washed with PBS, counted, and snap-frozen using liquid nitrogen. Crosslinking was performed using 2% formaldehyde according to the crosslinking protocol for cryopreserved cells provided by the Arima-HiC Kit user guide (A160134_v01). Following the manufacturer’s protocol, proximally ligated DNA was generated using Arima-HiC Kit (A510008). Hi-C library preparation was conducted using the KAPA HyperPrep Kit (Roche, 07962312001) and KAPA single-indexed adapter (Roche, 08005702001; index 4: TGACCA) according to Arima’s Library preparation user guide. Covaris S220 sonicator was used for DNA shearing. The library was quantified using the Qubit DNA HS assay (Life Technologies), and the library size was validated using DNA HS Bioanalyzer chips (Agilent). Paired-end sequencing was performed on the NovaSeq 6000 with 2x 100bp read length. The ligation sites of HiC were obtained using pre-set parameters in the Arima pipeline.

### Dot-plots

Two-dimensional (2D) dot-plots comparing chromosome assemblies from M. musculus (GRCm38-mm10), *M. spicilegus* (MUSP714_v2, GCA_03336285.2), *M. spretus*^61^ (SPRET_EiJ_v3, GCA_921997135.2) and *M. caroli*^62^ (CAROLI_EIJ_v1.1, GCA_900094665.2), and self-symmetry dot-plots showing internal duplications were generated from a custom Perl script^63^.

### Mapping t-complex inversion boundaries

To precisely map inversion boundaries, chromosomal regions spanning *t^w5^* inversion breakpoint junctions were estimated based on 2D dot-plots, repeat masked using default parameters on RepeatMasker^60^ , and compared to the mm10 chr17 sequence using BLAT^64^. Inversion boundaries were experimentally validated via PCR using primers flanking predicted *t*^w5^ inversion boundaries compatible with both *t^w5^* and mm10 chr17 sequences but only amplified from *t^w5^* genomic DNA (gDNA) (Extended Data Fig. 1B).

### Intrachromosomal identity, TE, and GC content

Sliding windows analyses were conducted to estimate percent intrachromosomal identity (50kb window, 1kb step), TE (200kb window, 10kb step), and GC content (100kb window, 10kb step). To calculate intrachromosomal identity, BLAST^64^ was used to compare all windows to the *t^w5^* assembly, and the top filtered results >10kb, not non-self-alignments. To calculate TE content, the *t^w5^* assembly was repeat masked using the default parameters of RepeatMasker^60^ , and percent SINE, LINE, and ERV elements were calculated for each window using a custom Perl script. Similarly, a custom Perl script calculated the percent GC content for each window.

### Sequence degeneration estimates

Single-copy sequence degeneration on the *t^w5^* was estimated via alignment to the first 50Mb of mm10 chr17 (excluding the centromere) using Minimap2 v.2.14^65^ and identifying regions with low *t^w5^*alignment coverage. Coverage was determined using bedtools^66^ , and the sum of non-overlapping, low-coverage base pairs was calculated using a custom Python script. Low *t^w5^* coverage regions were plotted relative to *wt* using a sliding window (100kb window, 10kb step).

### Copy number estimates

Copy number estimates were estimated using Illumina short-read sequencing data generated from homozygous *t^w5^*/*t^w5^* mESCs and previously published C57BL/6J^67^ (SRR7511358), *M. spretus*^61^ (ERR9880927), *M. caroli*^62^ (ERR133992), and *M. spicilgus*^68^ (SRR6356165) genomic DNA. We analyzed the short-read sequencing data via fastCN^69^, a pipeline for copy-number estimation based on read depth. Briefly, reads were mapped to the *t^w5^* or mm10 chr 17 assemblies in which RepeatMasker^60^, Tandem Repeat Finder^70^, and 50-mers with an occurrence greater than 50 were masked. Read depth was normalized to account for GC content, averaged in windows containing 3kb of unmasked positions, and converted to copy-number estimates based on defined single-copy sequence in the mm10 reference assembly.

### Illumina short-read sequencing

According to the manufacture protocol, mate-pair sequencing libraries were prepared from *t^w5^*/*t^w5^* mESC gDNA using the Illumina Nextera - MatePair kit and sequenced on an Illumina HiSeq2500

### Pachytene spermatocyte and round spermatid FACS and RNA-seq library preparation

A previously published protocol was followed to isolate pachytene spermatocytes (4N) and round spermatids (1N) from adult testes^71^. We disassociated cells from a pair of testes by enzymatic treatment with Collagenase type IA (Worthington Biochemical), DNaseI (Invitrogen), and Trypsin (Gibco). Cell suspensions were passed through 70µm and 40µm cell strainers and incubated with Hoechst 33342 (Thermo Fisher) for DNA content and propidium iodide for cell viability. Cell sorting was performed on the Moflo Astrios Cell Sorter. The purity of sorted cells was determined via fluorescence microscopy visual inspection of 100 cells morphology and γH2AX (Abcam, ab17722) immunofluorescence for pachytene spermatocytes or nuclear staining with DAPI for round spermatids. The purity of sorted cells used for subsequent RNAseq was >85%.

Cells sorted for RNA sequencing were sorted directly into TRIzol LS (Thermo Fisher Scientific) for total RNA extraction. RNA was treated with DNase using the RNeasy Mini Kit and the RNase Free DNase Set (QIAGEN) to remove any contaminating genomic DNA. RNA sequencing libraries were generated using samples with > 7 RIN scores and >5ng total RNA. mRNAseq libraries were generated using the NEBNExt Poly(A) mRNA Magnetic Isolation Module (NEB, E7490) and NEBNext UltraExpress RNA Library Prep Kit (NEB, E3330).

Pachytene spermatocyte mRNA-seq libraries were generated from three *wt/wt* and two age-matched *wt*/*t^w5^* biological replicates. Round spermatid mRNAseq libraries were generated from three biological replicates of *wt/wt* and *wt*/*t^w5^* males. All libraries were sequenced on a single Illumina NovaSeq Flowcell (∼25 million reads per sample on average). Alignments were performed with Kallisto^72^ using the RefSeq Index, and differential expression was estimated using DESeq2^73^ and plotted using ggplot2^74^.

### t^w5^ Gene Annotation

To generate gene annotations of the *t^w5^* sequence, we combined predicted gene annotations based on our sorted pachytene spermatocyte and round spermatid RNA-seq data with the mm10 RefSeq gene set. The RNA-seq reads were first normalized in Trinity ^75^ and then aligned to make second-strand cDNA predictions using *t^w5,^* the genome assembly as a guide, and an intron max length of 10kb. We aligned the Trinity-based and mm10 RefSeq-based cDNAs to the *t^w5^* assembly via BLAT^59^ using a minIdentity=92. We retained only the best hits with a minimum of 80% coverage using the Augustus ^76^ perl script filterPSL.pl and generated gff hints files for Augustus with the blat2hints.pl script. The Trinity-based RNA-seq and mm10 RefSeq gff hint files were combined and used in Augustus to annotate softmasked *t^w5^*sequence using Augustus with parameters –species=humans, softmasking=1, --UTR=on with configuration file extrinsic.M.RM.E.W.P.cfg. The Augustus gff outputs were converted to fasta files with gffread^77^. To identify *t^w5^* amplified gene families with *wt* orthologs, we compared the CDS of all annotated mm10 chr17 protein-coding genes within syntenic regions of estimated *t^w5^* amplicons (based on fastCN and self-symmetry triangular dot-plots). Genes with ≥2 *t^w5^*orthologs were considered ampliconic.

### Generation of an Amp2^tw5Δ^ and Amp2 ^wtΔ^ mouse lines

To generate mice with a ∼1.8Mb deletion of *t^w5^*-Amp2, loxP sites were sequentially integrated on the 5’ and 3’ flanking regions of Amp2 via CRISPR. sgRNAs were designed to target unique sequence flanking Amp2 on the *t^w5^,* selected based on the UCSC Genome Browser CRISPR Targets track scores, and synthesized by Synthego (Extended Data Table 5). sgRNA, Cas9 mRNA (IDT Alt R S.p. Cas9 Nuclease V3), and a single-stranded oligo donor carrying the loxP sequence (IDT; Extended Data Table 5) were mixed for microinjection at a concentration of 30, 50, and 10 ng/µl respectively. Pronuclear microinjection was performed on zygotes derived from the cross of *wt/t^w5^*males on a mixed C57B/6J and 129S4/SvJae background with C57B/6J/SJL F1 females. Heterozygous floxed mice (*Amp2^+/tw5-loxP^)* were mated against C57BL/6J *Ella-Cre* mice, resulting in a single mouse line carrying a deletion of the Amp2 region on the *t^w5^*(Fig. 4). Mice were genotyped by extracting DNA from tail or ear biopsy using NaOH extraction ^78^ and primers flanking the 5’ and 3’ loxP sites (Extended Data Table 5).

To generate mice with a ∼500kb deletion of *wt*-Amp2, CRISPR and sgRNA targeting the 5’ and 3’ flanking regions of *wt*-Amp2 and a homologous donor oligo were used. sgRNAs were designed to target unique sequence flanking Amp2 on the *wt* using the CRISPR/Cas9 10K Tracks in the UCSC genome browser (Extended Data Table 5). sgRNA, Cas9 mRNA (IDT Alt R S.p. Cas9 Nuclease V3), and a single-stranded oligo donor carry homology arms to the cut sites (IDT; Extended Data Table 5) were mixed for microinjection at a concentration of 30, 50, and 10 ng/µl respectively. Pronuclear microinjection was performed on zygotes derived from C57B/6J/SJL F1 males and females. Founder mice carrying the deletion were genotyped by extracting DNA from tail or ear biopsy using NaOH extraction^78^ and primers flanking the 5’ and 3’ sgRNA cut sites (Extended Data Fig. 7A, Extended Data Table 5).

### Transmission distortion assay

To assess the transmission frequency of *t^w5^or wt* alleles, we genotyped offspring resulting from *Amp2^+/tw5Δ^, Amp2^+/tw5-loxP^*, and *Amp2^w/wt^*^Δ^ males bred with CD1 females for *Amp2*^tw5Δ^, *wt* versus *t*^w5^, and *Amp2^wt^*^Δ^ alleles (Extended Data Table 5). At least three male mice were bred for each genotype to produce at least 100 offspring.

### RT-PCR

Total RNA was extracted using Trizol (Life Technologies) according to the manufacture protocol. Ten µg of total RNA was DNase treated using Turbo DNase (Life Technologies) and reverse transcribed using Superscript II (Invitrogen) using oligo (dT) primers following the manufacturer’s protocol. Intron-spanning primers were used to perform RT-PCR on adult testis cDNA for *Ppp1cb*, *Rnpepl1*, *Tmem181a-Rnpepl1,* and *Trim42*, a round spermatid specifically expressed gene and internal positive control.

### ddPCR

Droplet digital PCR (ddPCR) was conducted to estimate *Dynlt1* copy on the *t^w5^* and *wt* haplotypes experimentally. According to the manufacturer’s protocol, HMW DNA was extracted from snap-frozen spleen tissue using the MagAttract HMW DNA Kit (QIAGEN). Primers and TaqMan custom probes unique to *Dynlt1* and *Sox9,* a single copy internal control, were designed to amplify ∼150bp products (Integrated DNA Technologies; Extended Data Table 5). Products were amplified from two technical replicates for each DNA sample according to the Droplet Digital PCR Applications Guide (Bio-Rad) on the QX200 Droplet Generator and QX200 Droplet Reader and quantified using QuantaSoft Software.

### Defining Human and Mouse genome-wide amplicons

Genome-wide segmental duplications were collected from the UCSC Genome Table Browser for human (GRCh38/hg38) and mouse (GRcm38/mm10) genomes. Segmental duplications were filtered by size (>=10kb), identity (>=99%), and distance between duplications (<1.5Mb). We define this filtered set of segmental duplications as genome-wide amplicons. All UCSC Refseq annotated genes within amplicons were collected from the UCSC Genome Table Browser. To determine whether ampliconic genes are spermatid expressed, we aligned RNAseq data from sorted human (SRR1977587, SRR1977610, SRR1977618) and mouse (SRR1977639) spermatids^79^ to their respective RefSeq indices using Kallisto^72^, and defined expression as genes with >=10 tpm.

## Data Availability

The primary read files for the *t^w5^* sequence assembly, mate-pair sequencing and RNA sequencing, and *t^w5^* assembly are available through the National Center for Biotechnology Information (NCBI) (BioProject ID PRJNA1142398).

## Code Availability

Custom codes developed for data analysis and visualization are available upon request from the corresponding author.

## Acknowledgements

We want to acknowledge the Genome Technology Access Center at the McDonnell Genome Institute at Washington University School of Medicine for assistance with genomic analysis, the University of Michigan Flow Cytometry Shared Resource Laboratory for FACS, Advanced Genomics Core for RNA Sequencing and Transgenic Animal Model Core for generating mouse deletion lines. We are grateful to Christian Rohrandt for his invaluable input on nanopore data analysis and preprocessing. We thank the MPIMG IT and SeqCore facility for Mate-Pair and PacBio library preparation and sequencing, particularly for the support from Thomas Kreitler. We thank M. Meisler for sharing *Ella-Cre* mice, T. Keane for sharing a draft chr17 *M. spretus* assembly, D. Tautz, L. Odenthal-Hesse, and K. Ulrich for sharing *M. spicilegus* and *M. spretus* tissue and a *M. spicilegus* chr17 assembly, and D.W. Bellott for technical advice. We thank M. Arlt, D.W. Bellott, E. Clowney, J. Kidd, A. Lawson, J. Moran, T. Wilson, and Y. Yamashita for their comments.

## Author contributions

C.M.S. performed overall project coordination, experimental and computational analyses, and figure generation. G.W. performed long-read (Oxford Nanopore and PacBio HiFi) data processing, integrated long-read and Hi-C genome assemblies, and manually curated the *t^w5^* region. L.Z. performed ddPCR and genotyping for transmission distortion assays. B.B. prepared DNA for ultra-long Nanopore and PacBio HiFi sequencing. H.B. generated the *t^w5^/t^w5^*mouse embryonic stem cell (mESC) line. P.T. conducted Hi-C analysis from *t^w5^/t^w5^*mESCs. F.J.M., B.G.H., and J.L.M. provided project leadership and coordination. C.M.S and J.L.M. wrote the manuscript with contributions from other authors.

## Ethics Declarations

### Animal Use

All animal experiments were conducted in compliance with IACUC (PRO00011089).

