## Supplementary Information for "Acquisition of ampliconic sequences marks a selfish mouse *t*-haplotype"

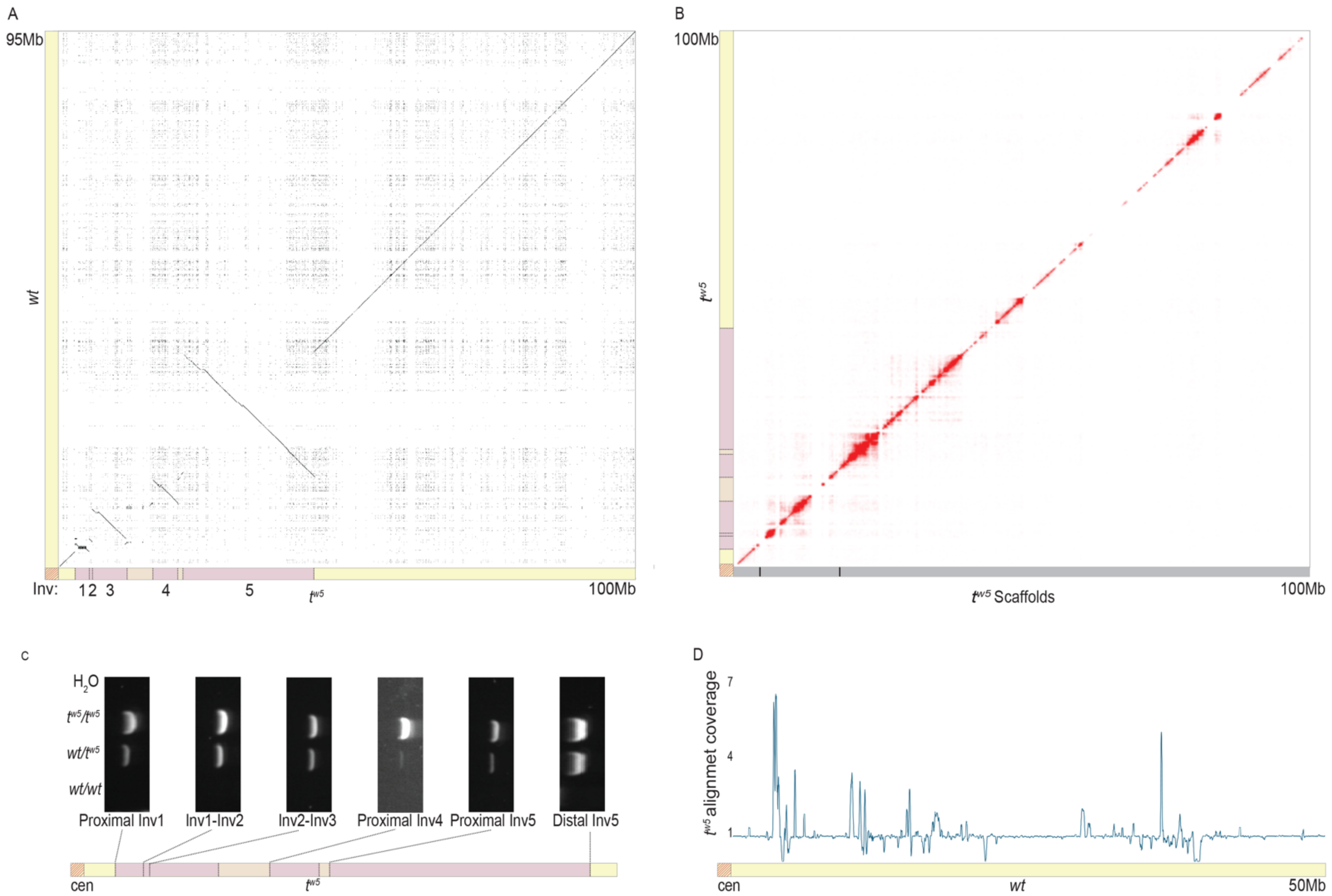

**Figure 1. Assembly of the mouse chromosome 17 $t^{w5}$**

**A.** 2-D dot-plot comparison of the $t^{w5}$ sequence assembly (X-axis) and the mm10 reference chromosome 17 (*wt;* Y-axis). Each dot represents 100% nucleotide identity of a 250bp DNA segment between the two assemblies, with long stretches of similar sequence represented as lines. Along the $t^{w5}$ chromosome, dark purple shading represents inverted regions, and light purple shading represents non-inverted regions. Dotted lines represent inversion junctions. Red diagonal lines represent the centromere. Five inversions between the $t^{w5}$ and *wt* chromosomes are numbered below. **B**. Contact map of Hi-C data generated from $t^{w5}/t^{w5}$ mESCs and mapped to $t^{w5}$ assembly scaffolds (grey bars). Three non-overlapping hybrid scaffolds (4Mb, 14Mb, and 81Mb) are represented by a grey line with black bars denoting physical gaps between scaffolds. **C.** PCR validation of $t^{w5}$ inversion junctions. **D.** A coverage plot of the $t^{w5}$ assembly aligned to the first 50Mb of *wt* (mm10 chr17) using a sliding window (100kb window, 10kb step).

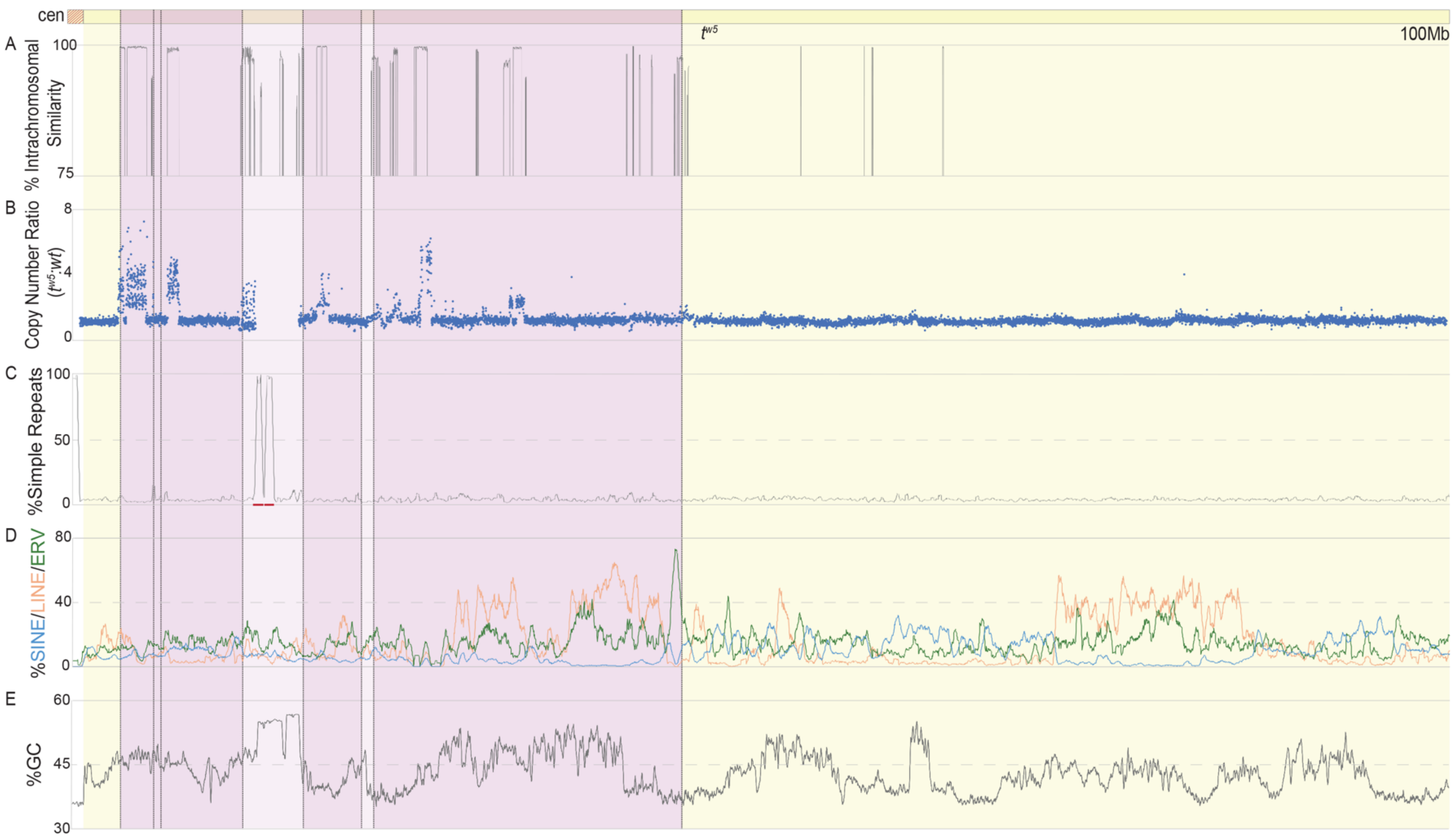

**Figure 2. $t^{w5}$ sequence content**

**A.** Intrachromosomal sequence similarity using a 50-kb sliding window and 1-kb steps. Each window was compared to all other windows, excluding self-comparisons and transposable elements. **B.** $t^{w5}$ copy number estimates relative to wt (Y-axis) based upon read-depth analysis of whole-genome short-read sequencing data from $t^{w5}/t^{w5}$ mESCs and C57BL/6J adult mice mapped to $t^{w5}$ chr17 (X-axis). Each dot represents copy number estimates in a 3kb window of unique sequence. Copy number ratios >1 represent sequence of increased copy number on the $t^{w5}$. **C.** Simple nucleotide repeat content as a percentage of nucleotides in a 200-kb sliding window with 1-kb steps. Two blocks of $t^{w5}$-specific simple repeats are underlined in red. **D.** LINE, SINE, and ERV content as a percentage of nucleotides in a 200-kb sliding window with 1-kb steps. **E.** G + C content (%) calculated in a 100-kb sliding window with 1-kb steps.

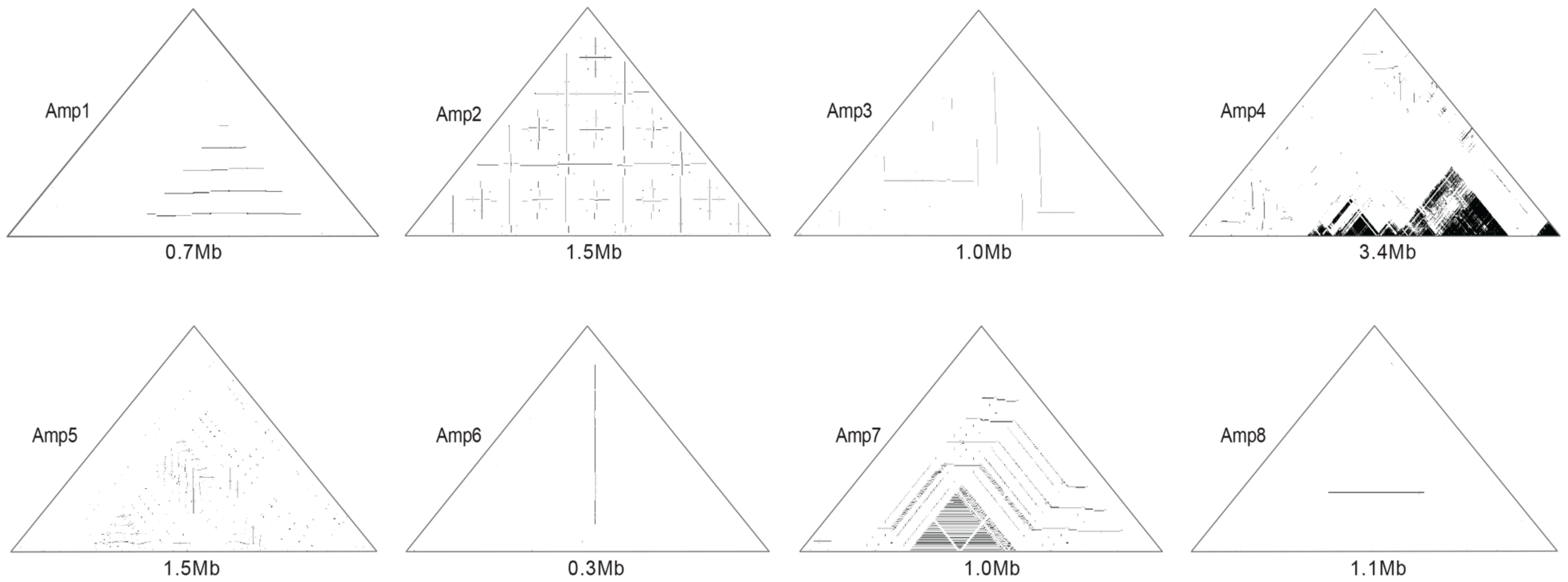

**Figure 3. Eight $t^{w5}$ ampliconic regions**

Self-symmetry dot-plots highlighting palindromic (vertical lines) and tandem (horizontal lines) segmental duplications within $t^{w5}$ Amp1-8. Each dot represents a 100bp window of 100% nucleotide identity. The X-axis is the ampliconic sequence, 5’-3’.

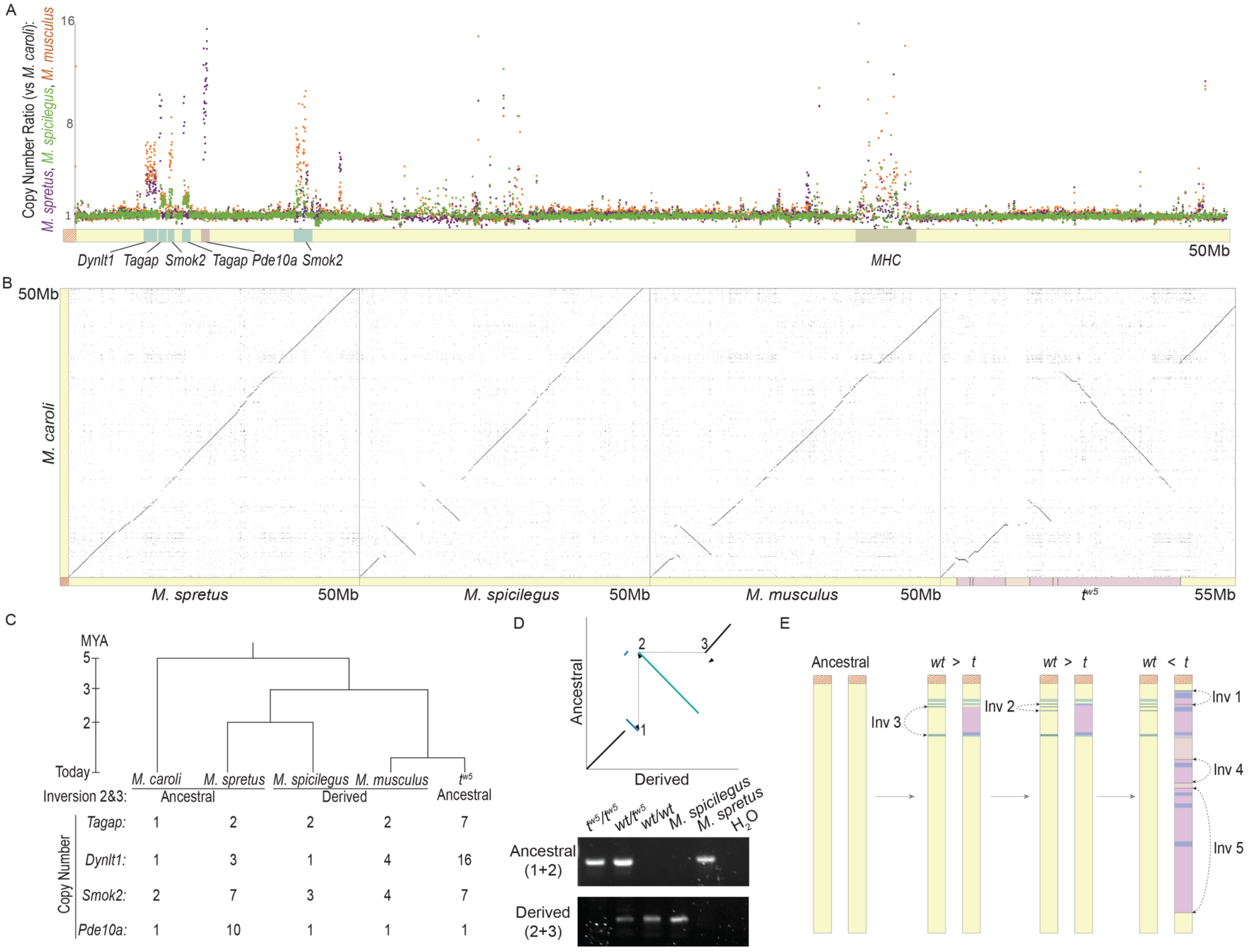
A
Copy Number Ratio (vs M. caroli):
M. spretus, M. spicilegus, M. musculus
16
8
1
Dynlt1 Tagap Smok2 Tagap Pde10a Smok2
MHC
50Mb
B
50Mb
M. caroli
M. spretus
50Mb
M. spicilegus
50Mb
M. musculus
50Mb
tw5
55Mb
C
MYA
5
3
2
Today
M. caroli
M. spretus
M. spicilegus
M. musculus
tw5
Inversion 2&3:
Ancestral
Derived
Ancestral
Copy Number
Tagap: 1 2 2 2 7
Dynlt1: 1 3 1 4 16
Smok2: 2 7 3 4 7
Pde10a: 1 10 1 1 1
D
Ancestral
Derived
1
2
3
tw5/tw5
wt/tw5
wt/wt
M. spicilegus
M. spretus
H2O
Ancestral (1+2)
Derived (2+3)
E
Ancestral
wt > t
wt > t
wt < t
Inv 3
Inv 2
Inv 1
Inv 4
Inv 5

**Figure 4. Conserved sequence amplification is genetically linked to the first *t*-complex inversions**
**A.** Copy number estimates across the first ~50Mb of *M. spretus*, *M. musculus (*X-axis*),* and *M. spicilegus* chr17 relative to *M. caroli* (Y-axis). Each dot represents a copy number estimate across a 3kb window of a unique sequence. Conserved amplicons are shaded in blue on the X-axis. Unique amplification in *M. spretus* is shaded in blue, and the *MHC* locus is shaded in grey. Gene families are below the X-axis. **B.** 2-D dot-plot comparison of the first ~50-55Mb of *M. spretus, M. spicilegus, M. musculus and* $t^{w5}$ chr17 (X-axis) and the first 50Mb of *M. caroli* chr 17 (Y-axis). Each dot represents 100% nucleotide identity of 100 nucleotides, where diagonal lines indicate syntenic blocks of sequence. **C.** A schematic phylogenetic tree of *Mus* chr17 with the inversion 2 and 3 orientation (ancestral or derived) and copy numbers for ampliconic gene families below each species. **D.** A schematic 2-D dot-plot (top) highlighting the ancestral (vertical dotted line) versus derived (horizontal dotted line) inversion junctions with black arrows denoting primer positions for PCR validation (bottom) of ancestral versus derived junctions from $t^{w5}/t^{w5}$, *wt*/$t^{w5}$, *wt/wt*, *M. spicilegus,* and *M. spretus* gDNA. **E.** A schematic representation of step-wise inversions (Inv) and amplicon acquisition on *wt* and *t* haplotypes. Hypothesized selfish transmission of the *wt* vs. *t* over evolutionary time is indicated above each chromosome by ">" or "<." Black dotted arrows denote inversions, and blue-shaded regions denote amplicons. Horizontal dotted lines represent inversion junctions. Along the *t* chromosome, dark purple shading represents inverted regions, and light purple shading represents non-inverted regions between *t* and *wt*. Red diagonal lines represent the centromere.

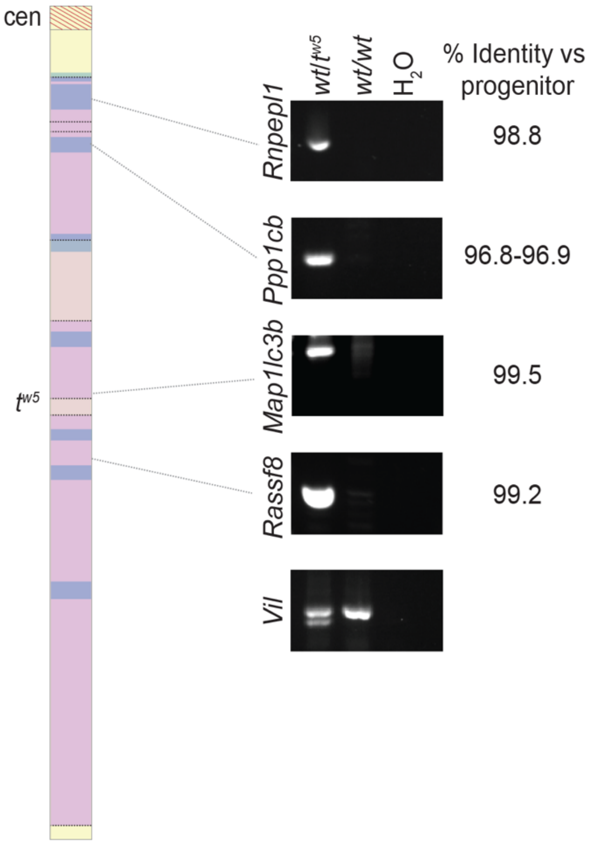

**Figure 5. Acquisition of four $t^{w5}$ retrogenes**
PCR validation of retrogene insertions using $t^{w5\text{-}RTG}$ specific primers, their locations on the $t^{w5}$ schematic (dotted grey line), and percent sequence identity to their progenitor gene copies.

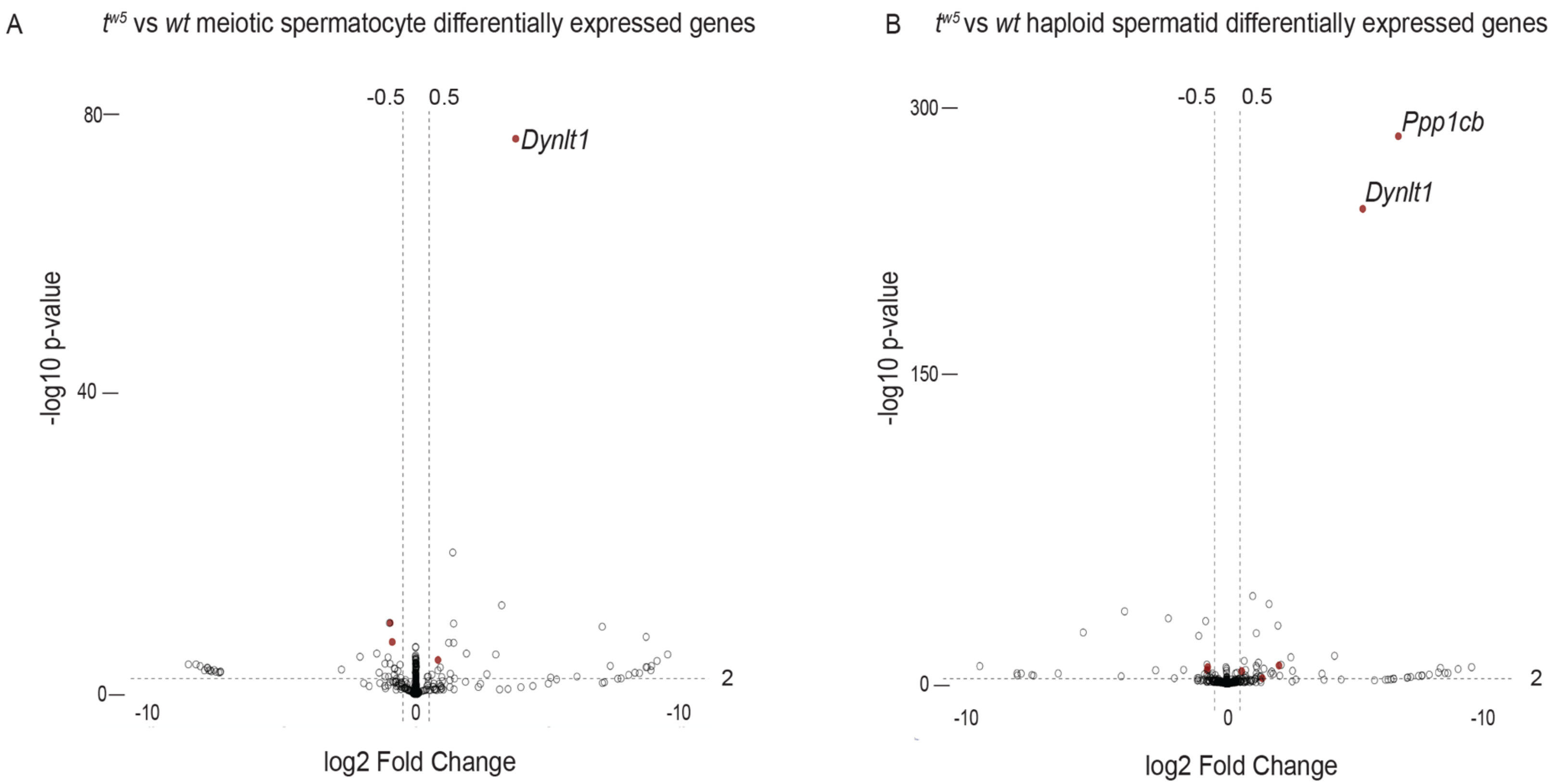

**Figure 6. Differential expression of $t^{w5}$ amplicon gene families in meiotic spermatocytes and haploid spermatids.** Volcano plots of differentially expressed genes (-log10 p-value > 0.01; log2 fold change > 0.5) in $t^{w5}$/*wt* versus *wt/wt* meiotic spermatocytes (A) and haploid spermatids (B). Filled red dots represent $t^{w5}$ ampliconic gene families with differential expression.

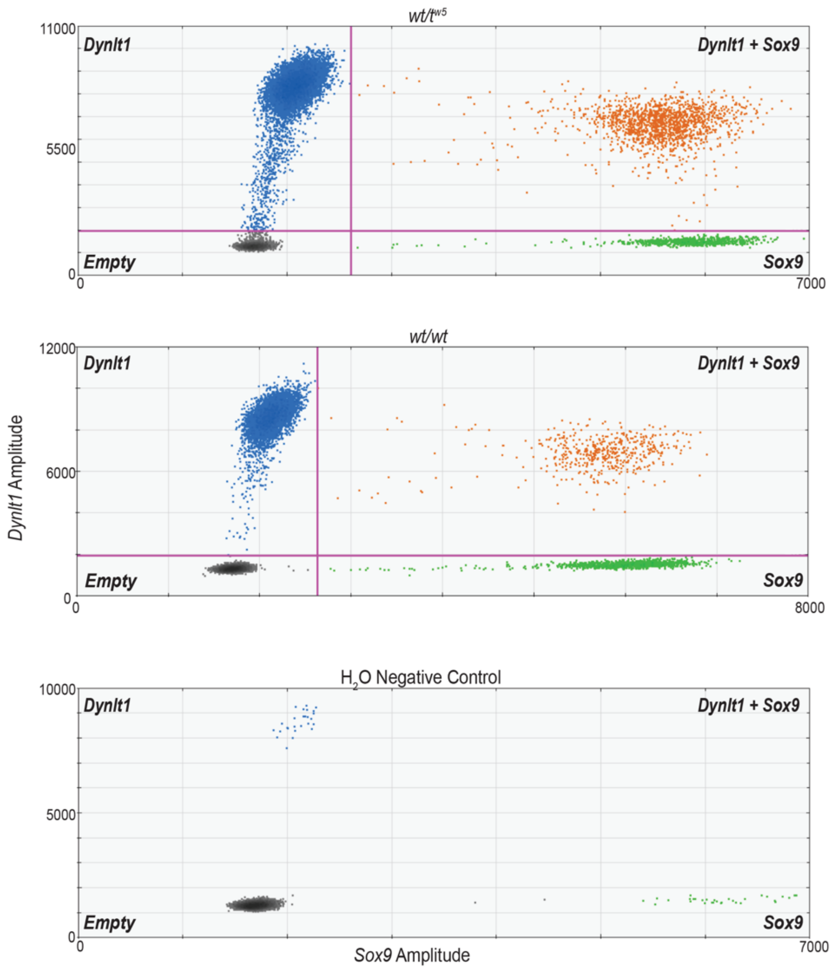

| | Concentration | Ratio | $t^{w5}$ CN |
|---|---|---|---|
| ***Dynlt1*** | 1356 | 20:2 | 16 |
| ***Sox9*** | 137 | | |

| | Concentration | Ratio | *wt* CN |
|---|---|---|---|
| ***Dynlt1*** | 428 | 9:2 | 4.5 |
| ***Sox9*** | 95 | | |

**Figure 7. Droplet digital PCR (ddPCR) estimation of *Dynlt1*$^{tw5}$ versus *Dynlt1*$^{wt}$ copy number**

ddPCR droplet plots showing the distribution of target DNA into droplets. X-axes are fluorescence amplitude corresponding to *Sox9*, a single-copy internal control, and the Y-axes are fluorescence amplitude corresponding to *Dynlt1*. The Top left quadrant contains *Dynlt1* positive droplets (blue), and the bottom left quadrant contains *Sox9* positive droplets (green). The bottom left quadrant contains negative droplets (grey), and the top right contains droplets positive for *Dynlt1* and *Sox9* (orange). Tables corresponding to droplet plots include the estimated concentration of *Dynlt1* and *Sox9* in each sample, the ratio of *Dynlt1:Sox9* concentration, and the estimated copy number for each allele. ddPCR was conducted using $t^{w5}$/*wt* (top) and *wt/wt (middle)* gDNA with water as a negative control (bottom).

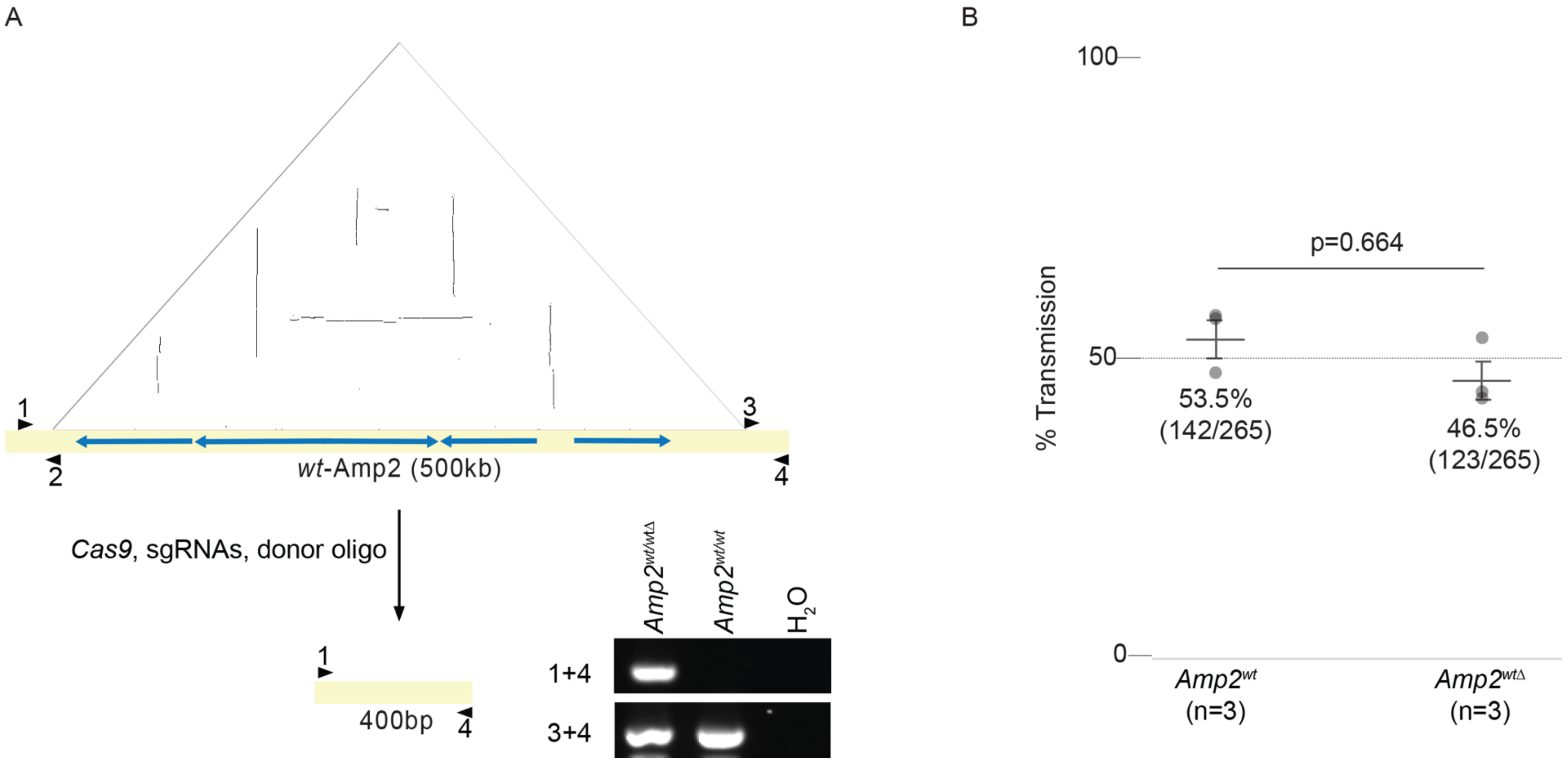

**Figure 8. Deletion of *wt*-Amp2 does not influence transmission ratio**

**A.** (Top) Self-symmetry dot plot highlighting palindromic (vertical lines) and tandem (horizontal lines) segmental duplications within *wt*-Amp2. Each dot represents a 100bp window of 100% nucleotide identity. The X-axis is the ampliconic sequence, 5'-3', with a blue arrow representing segmental duplications. Black arrows with numbers denote primer positions flanking sgRNA target sites used for genotyping. (Bottom) A schematic of the *wt*-Amp2 region targeted for deletion via sgRNA guided Cas9 and a donor oligo and PCR validation of the $Amp2^{wt\Delta}$ from genomic DNA. **B.** $Amp2^{wt/tw5\Delta}$ (n=3) males were mated against WT females, and progeny were genotyped for $Amp2^{wt\Delta}$. Average percent transmission (horizontal line) and number of $Amp2^{wt}$ or $Amp2^{wt\Delta}$ alleles/total progeny genotyped in parentheses are shown. Vertical lines represent standard error. The p-value was calculated using a generalized linear mixed model. Each dot represents the percent allele transmission from a single stud male.

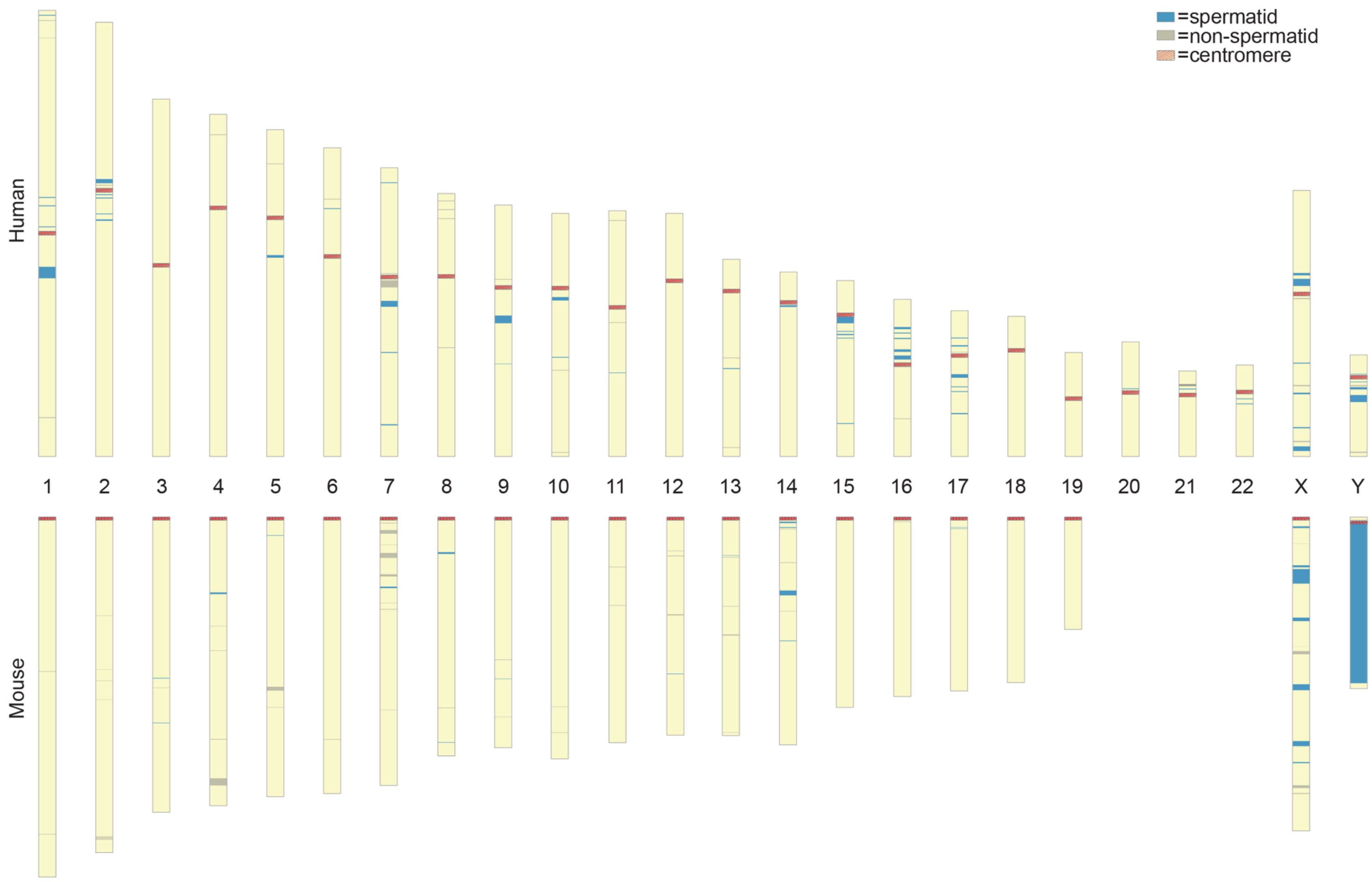

**Figure 9. Amplicons with spermatid-expressed genes across human and mouse genomes.**
Chromatograms of human (hg38) and mouse (mm10) genomes with amplicons containing spermatid expressed (blue) or non-spermatid expressed (grey) gene families.

**Table 1. Inversion boundaries**

| Inversion | $t^{w5}$ start | $t^{w5}$ end | Size (kb) | *wt* start* | *wt* end* |
|---|---|---|---|---|---|
| 1 | 3729394 | 6186130 | 2457.7 | 5747331 | 6957919 |
| 2 | 6185539 | 6690983 | 505.4 | 6957919 | 7435308 |
| 3 | 6690248 | 13208519 | 6518.3 | 7435308 | 13016510 |
| 4 | 17088738 | 21377882 | 4289.1 | 14068229 | 18051507 |
| 5 | 22371805 | 44778705 | 22106.9 | 19401000 | 40049964 |

*Wild-type (*wt*) coordinates are from the mm10 chr17 reference sequence

**Table 2. Inversion $t^{w5}$ junction regions**

| Inversion (Inv) junction | Inversion junction $t^{w5}$ position | PCR product size (bp) |
|---|---|---|
| Proximal Inv1 | 3729394-3730605 | 1211 |
| Inv1-Inv2 | 6185539-6186130 | 591 |
| Inv2-Inv3 | 6690248-6690983 | 735 |
| Proximal Inv4 | 17088738-17089744 | 1006 |
| Proximal Inv5 | 22371805-22372812 | 1007 |
| Distal Inv5 | 44778196-44778705 | 509 |

**Table 3. *wt* vs. $t^{w5}$ sequence differences**

| | ***wt**** | **$t^{w5}$** | **%Δ** | **bpΔ** |
|---|---|---|---|---|
| Total Bases (Mb) | 34.28 | 41.05 | +19.75 | +6.77 |
| Satellite Repeats (Mb) | 0.85 | 3.73 | +341.02 | +2.88 |
| TEs (Mb) | 14.07 | 16.18 | +15.01 | +2.11 |
| Low $t^{w5}$ Coverage (Mb) | - | - | - | -3.80 |

*Wild-type (*wt*) sequence estimates are from the mm10 chr17 reference sequence

**Table 4. $t^{w5}$ Amplicons**

| | $t^{w5}$ position (Mb) | Repeat unit size (kb) | Total ampliconic sequence (kb) | Gene families | Copy number |
|---|---|---|---|---|---|
| Amp1 | 3.7-4.1 | 15.3-139.2 | 357.1 | ***Tagap*** | 7 |
| | | | | ***Rsph3*** | 5 |
| Amp2 | 4.3-5.7 | 10.1-244.0 | 1401.8 | ***Dynlt1*** | 13 |
| | | | | ***Tmem181a*** | 12 |
| | | | | ***Rnpepl1***$^{tw5\text{-}RTG}$ | 4 |
| | | | | ***Tmem181a-Rnpepl1***$^{tw5\text{-}RTG}$ | 4 |
| | | | | ***Sytl3*** | 6(5) |
| | | | | *Tulp4* | 7(6) |
| Amp3 | 7.2-8.0 | 25.4-26.1 | 818.1 | ***Slc22a3*** | 4 |
| | | | | ***Ppp1cb***$^{tw5\text{-}RTG}$ | 4 |
| | | | | *Plg* | 3 |
| Amp4 | 12.6-15.5 | 10.1-64.9 | 2964.5 | ***Smok2*** | 7 |
| | | | | ***Rps6ka2*** | 3 |
| | | | | ***Tcp10c*** | 4 |
| Amp5 | 18.0-18.7 | 30.2-71.9 | 706.6 | *Zfp960* | ~8 |
| Amp6 | 23.5-23.8 | 103.2 | 249.5 | *Olfr136* | 2 |
| | | | | *Esp36* | 6(5) |
| | | | | *Exp34* | 5(4) |
| Amp7 | 25.0-25.9 | 13.4-126.9 | 921.9 | ***Ppp1r11*** | 5 |
| | | | | ***Gabbr1*** | 2(1) |
| | | | | *Znrd1* | 5 |
| | | | | *Znrd1as* | 5 |
| | | | | *H2-M5* | 5(4) |
| | | | | *Olfr90* | 2 |
| Amp8 | 32.2-32.7 | 129.8 | 568.2 | ***Dnah8*** | 2 |
| | | | | ***Glo1*** | 2(1) |
| | | | | ***Rsph1*** | 2(1) |
| | | | | ***Slc32a1*** | 2(1) |
| | | | | *Tff1* | 2(1) |
| | | | | *Tmprss3* | 2(1) |
| | | | | *Ubash3a* | 2 |

() = partial copies; ~ = approximate copy number; ***Spermatid expressed***; *Not spermatid expressed*

**Table 5. List of oligos and gRNA**

**Oligos for Genotyping**

| Name | Sequence |
|---|---|
| Vil_F | TCATGGACCAACACAAGCTC |
| Vil_R | CACAAAACTGAAATCTCCCTCTC |
| HB_F | GAGTGACCTGCATGCCCACAAGCTGTG |
| HB_R | GACCTGTGGAGACAGGAAGGGTCAGTG |
| tw5_Amp2_5'F | ACGAGTCCAAAAGAACACGC |
| tw5_Amp2_3'R | GAGTCTCAGTCCCTACAGCA |
| tw5_Amp2_5'loxP_R | GCCCAGGTGTTAGCTTGAGA |
| tw5_Amp2_3'loxP_F | CGATTCCAGACCCCAACAAC |

**Oligos for Retrogene Validation (PCR and RT-PCR)**

| Name | Sequence |
|---|---|
| Ppp1cb_F | AAACGCCTAGCCAACACCTA |
| Ppp1cb_R | TGAGTCTGTCCACGTTCAGC |
| Rnpepl1_F | TCTCTGTATCGTGTGGAGCC |
| Rnpepl1_R | GTGCCAGTGCCGTGTGGGCA |
| Map1lc3b_F | ATCACTGGGATCTTGGTGGG |
| Map1lc3b_R | GGGTAGGGGAACTTGAGCTT |
| Rassf8_F | ACACAGCTGCTTGTAAGTGC |
| Rassf8_R | CCCTTGGGAGCTCATTAGGA |
| Tmem181a-Rnpepl1_F | AGTGAAGGTCGATGGTGTGT |
| Tmem181a-Rnpepl1_R | CTAGGCAGACACATTGACGT |
| Trim42_F | GAAGCATCGTCACCTCCTCT |
| Trim42_R | CTTCTCGCATAGGCTGTGGT |

**Oligos for ddPCR**

| Name | Sequence |
|---|---|
| Dynlt1_exon3_F | GAGTTGGCTCAAAGTCTG |
| Dynlt1_exon3_R | CCCCTAGGCTATAGAAAG |
| Dynlt1_exon3_Probe | TGGTTGACTTTGCTGTGCTGGT |
| Sox9_exon1_F | GAGCCGGATCTGAAGAAGGA |
| Sox9_exon1_R | CTTGACGTGTGGCTTGTTCT |
| Sox9_exon1_Probe | TGCTGAAGGGCTACGACTGGA |

**Oligos for $t^{w5}$ inversion junction PCR**

| Name | Sequence |
|---|---|
| Proximal Inv1_F | CCTGTTCAGGTCGGTTCTGT |
| Proximal Inv1_R | GGGCTGTAGGCAATCCTCTA |
| Inv1-Inv2_F | TGTTCCCAAAAGGGTCTCTG |
| Inv1-Inv2_R | GGCCTCACTGTGGAAACAAT |
| Inv2-Inv3_F | TCCTCTGACCTAAGTGGAACC |
| Inv2-Inv3_R | TCATGCATGCAAATCCTTGT |
| Proximal Inv4_F | TGGGCATAAGTGACAAGAAG |
| Proximal Inv4_R | ATTTGGTTGTTGGGTGGTGT |
| Proximal Inv5_F | TGCTATATGTTTTGAGAACAAAGTGA |
| Proximal Inv5_R | GCATAGCATGTATGGACTCAACA |
| Distal Inv5_F | CCATCCCAACCACACTTTTT |
| Distal Inv5_R | AAGTTGCTGTGCTTGTGCAG |

**Oligos to generate *Amp2* $^{tw5\Delta}$ *and Amp2* $^{wt\Delta}$**

| Name | Sequence |
|---|---|
| Amp2 $^{tw5}$_5'_donor | GACACGAGTCCAAAAGAACACGCTGGACCCGACTTTCGAGGAAACCTTGAAGGTA<br>CGTGCCATAACTTCGTATAATGTATGCTATACGAAGTTATAGGCGGGAAAACATT<br>CCTGAGTCCTCCCCACCTGGGCTAGGTCTAGGGTCAGAGTCCAGC |
| Amp2 $^{tw5}$_3'_donor | CGAGACTTTGTCAGCTACCCCTCAGCGGGCAATGAACGGCTGCACTGCACCATGA<br>AGCGCAATAACTTCGTATAATGTATGCTATACGAAGTTATCAGAGGATGACCCAG<br>AGGTCGGCGGGCCCTGCTACACACTCTACTTGGAGTACCTGGGAG |
| Amp2 $^{wt}$_donor | CCAAATCTAAAGCCTCAAGAGGTGCCTCAGAAGTACCAAAAAAAAAAACAAAAAT<br>AAAATAAAATAAAATACTCATTGGCACAAAATGAATGATCAAGGTATGCCTCTTT<br>GTAATGCTTGAGTTAGTTACACAGAACTGCAGACACCTGTACATTTCTTCAGGGT<br>TCTGTCAGTCAGTACAGGAGGCAGACGCCAGCCTT |
| Amp2 $^{tw5}$_5'_gRNA | CCTTGAAGGTACGTGCCAGGCGG |
| Amp2 $^{tw5}$_3'_gRNA | CTGCACCATGAAGCGCACAGAGG |
| Amp2 $^{wt}$_5'_gRNA | AUGAAUGAUCAAGGUUACGG |
| Amp2 $^{wt}$_3'_gRNA | AGCAUUACAAAGAGGCAUGG |
